# Molecular profiling of glioblastoma-derived extracellular vesicles identifies small nucleolar RNAs as candidate liquid biomarkers for radiation- induced senescence

**DOI:** 10.64898/2025.12.12.693499

**Authors:** Valerie DeLuca, Nathaniel Hansen, Priya Digumarti, Nanyun Tang, Karen Fink, George Snipes, Patrick Pirrotte, Michael Berens

## Abstract

Radiation-induced senescence (RIS) in glioblastoma (GBM) is an undesirable cell fate that reduces tumor cell death and supports resistance and outgrowth. While senescence-targeting drugs are promising adjuvants, their clinical application will require proper patient selection based on post-treatment RIS burden. Current methods to evaluate senescence, however, are tissue-based, and given GBM’s difficult anatomical location, post-treatment biopsies are impractical. Innovative and less invasive biomarkers for RIS are urgently needed. To this end, we aimed to identify candidate extracellular vesicle (EV) liquid biomarkers for RIS by profiling senescence-associated cargo changes within GBM EVs. Using a panel of GBM patient-derived cell cultures, we show that RIS is the primary functional state following radiation exposure and is associated with significant alterations in the cargo of senescent-derived EVs (senEVs). In particular, senEV transcriptomes have an increased abundance of senescence-associated RNA genes and gene sets. Most striking, however, was that senEVs are most differentiated by the significant enrichment of a panel of snoRNAs. This signature was conserved in 4/5 GBM models of RIS and was validated by qRT-PCR. We further confirmed snoRNA enrichment in the senEVs of a breast cancer cell line, as well as the lack of snoRNA enrichment following senescence-independent drug exposure. Analysis by mass spectrometry revealed that snoRNAs are likely co-packaged with their associating proteins, as senEVs had concurrent increases in these binding partners. We examined whether packaging is associated with nucleolar stress during RIS, but found that upregulation in senEVs is likely due to a tightly controlled cellular abundance rather than nucleoli fragmentation and release of nucleolar components to the cytoplasm. Finally, using a preliminary cohort of longitudinal plasma samples from four GBM patients, we determined the feasibility of detecting senescence-associated and snoRNA species in the extracellular vesicles of patient biofluids. Of interest, post-treatment EVs had increased senescence-associated RNAs like *CDKN2B* and *GLB1* and the snoRNA *SNORA49*. Altogether, this data suggests that senEV RNA species, and particularly snoRNAs, are a promising analyte for RIS-biomarker development. With further study, this work may open avenues for a companion diagnostic for senotherapeutics.

## 1. INTRODUCTION

Glioblastoma treatment options have failed to advance within the last decade, and barring enrollment to a clinical trial, patients are generally constrained to surgical resection, radiation (IR), and temozolomide (TMZ) chemotherapy^1,2^. There is a dire need for new therapeutics, but the drastic heterogeneity, stemness, and physical location of GBM behind the blood-brain-barrier all pose hindrances to the drug development process^3,4^. While novel first-line therapies are in development and hold promise^2^, an additional alternative is to exploit vulnerabilities induced by standard of care (SOC). This includes leveraging druggable persister states^5^, such as therapy-induced senescence (TIS).

TIS—or when promoted by radiation, RIS—is characterized by a prolonged growth arrest, cytoskeletal changes, enhanced lysosomal biogenesis, metabolic dysregulation, epigenetic rewiring (senescence-associated-heterochromatic-foci formation, SAHF), and an enhanced and altered secretome (senescence associated secretory phenotype, SASP)^6^. There are multiple cell-intrinsic and extrinsic consequences of TIS. Intrinsically, TIS tumor cells may serve as a reservoir for recurrent disease due to the potential for proliferative recovery^7^. While this is in contrast to the historical definition of senescence as an irreversible growth arrest, proliferative recovery from TIS has been documented in lung, breast, AML, prostate, and GBM cells^7–12^. Extrinsically, the SASP from both senescent tumor and senescent host cells can modulate the tumor microenvironment and contribute to immune evasion, chemotherapy-associated side effects, and tumor recurrence^13–17^. The ablation or modulation of senescent cells by senotherapeutics has been shown to enhance *in vitro* and *in vivo* drug response and longevity in various tumor types^11,18–21^. This includes promising preclinical results demonstrating the potency and tumor-control capacity of senolytic therapies in GBM^14,22–25^.

Senotherapeutic potency, by definition, is dependent upon having an appreciable senescent burden. For example, senolytics have minimal toxicity to proliferating cells and sensitivity following exposure to senescence-stimuli is time sensitive. For example, Rahman *et al.* found that potency to the senolytic ABT-263 increases with time as GBM cells enter senescence following radiation exposure^23^, while Saleh *et al.* demonstrated that potency decreases with time as lung cancer cells recover from etoposide-induced senescence^20^. TIS is likely also heterogeneous between patients, as evaluations of breast and lung cancer have shown that neoadjuvant chemotherapy induces TIS in ∼40% of patients^8,26,27^. However, the prevalence of senescence and the incidence promoted by SOC in GBM patients are not well understood. While early studies in resected tissue support a minimal pre-existing senescent fraction in treatment-naive tumors (<10% in any given section)^17,28^, studies of TIS have been hampered by the inaccessibility of post-treatment tissue prior to recurrence. Further, biopsies are apt to miss TIS in regions that are unresectable, such as adjacent normal brain tissue. A tissue-independent assay for senescence in GBM patients might therefore i) enable a better understanding of the heterogeneous presentation and clinical consequences of TIS/RIS and ii) serve as a companion diagnostic for senotherapeutics by identifying patients with an increased senescent burden post-SOC.

Extracellular vesicles (EVs) have gained increased interest as a liquid biopsy for diagnosis and disease monitoring, including in GBM^29–31^. EVs are membranous particles secreted by both tumor and host cells that contain RNA, DNA, and protein^32^. EVs from senescent cells (senEVs) *in vitro* have been shown to contain unique cargo^33–36^, and EV protein cargo has been explored as a potential biomarker for senescent fibroblasts and Warner’s syndrome^37^. However, to our knowledge, it has yet to be established if and how EVs may report on RIS in GBM, where markers may differ from noncancerous cell types.

Here, we investigate the elaboration of senEVs and their cargo in senescent GBM cells to qualify the potential of an EV-based TIS/RIS bioassay. Our results show that senescence-associated RNA species and snoRNAs are significantly enriched in senEVs. Importantly, snoRNA enrichment was conserved across 4/5 GBM models, which is promising for a heterogenous phenotype, in a heterogenous cancer. Using longitudinal patient samples obtained at surgical resection and following completion of SOC, we further show the feasible detection of these species in patient plasma EVs. Finally, we show that enrichment of snoRNAs in senEVs is likely not a consequence of nucleolar stress/damage during RIS but rather related to homeostasis of snoRNA transcription and release. We ultimately conclude that radiation-induced senescence in GBM results in the elaboration of EVs that are differentiated by a signature comprised of senescence-associated RNA species and small nucleolar RNA. These EV-transcriptome changes warrant additional investigation and development as a more targeted biomarker/bioassay for RIS.

## 2. MATERIALS AND METHODS

### 2.1. Cell culture and treatments

GBM patient-derived cultures (GBM6, GBM43, GBM102, and GBM120) were acquired from Mayo Clinic^38^. A172, H460, and MDA-MB-231 cells were acquired from ATCC. Cell line characteristics are summarized in Table S1. Media components are listed in Table S2. Patient-derived cultures were grown and generally tested as spheroids, but plates or slides were pre-coated with laminin (Sigma Aldrich, L2020) for microscopy endpoints to aid visualization. All tumor lines were authenticated by STR and tested negative for mycoplasma. Irradiation was performed with the X-Rad320 system (Precision X-ray, Inc), housed at University of Arizona, Phoenix Biomedical Campus. Temzolomide (TMZ) was suspended in DMSO and cells treated for 72 hours. Time of treatment is considered day/hour 0. Controls are generally harvested on day 3, when they are in logarithmic and viable growth, and treated cells on day 7, unless otherwise indicated.

### 2.3. Drug dose response

Cells were plated in triplicate wells of a 96-well plate, allowed to adhere overnight, and then treated. For radiation, shields were used to ensure appropriate dosing. Relative viability was evaluated with CellTiter-Glo (Promega, G7570).

### 2.4. Live cell imaging

Live cell imaging was performed in 96-well plates on the CellCyte X^TM^ (Cytena) at 10X. Analysis was performed by the CellCyte software, using the cell confluency program. Conditions for each independent experiment were analyzed in duplicate, at minimum.

### 2.5. SA-β-galactosidase staining

Histochemical staining by XGal (Invitrogen, 15520018) was performed as described previously^39^. Cells were imaged on a Keyence BZ-X800 at 20X and white balancing/brightening was performed in the Keyence analysis software. All shown images are representative fields from at least three independent experiments unless otherwise stated. Quantification was performed in ImageJ by converting images to 8-bit and then thresholding and measuring percent of image area for thresholds capturing blue staining. This percentage was then normalized against percent area of the cells in the image, to adjust for variable confluency.

### 2.6. Western Blotting

Cells or EVs were lysed in 1X RIPA (Millipore, 20-188) with cOmplete protease inhibitor cocktail (Roche, 4693116001) and phosphatase inhibitor cocktails I and II. Protein quantification was performed with the Pierce (Micro) BCA Protein Assay Kit (Thermo Scientific, 23225 and 23235). Equivalent protein quantities were separated by SDS-PAGE with NuPAGE 4-12% Bis-Tris gels (Invitrogen, NP0335) and then transferred to 0.45 micron polyvinylidene membranes (Invitrogen, LC2005). Membranes were blocked with 5% BSA and probed overnight at 4℃ for primary antibodies and probed for 2 hours at room temperature for secondary antibodies. Antibody information can be found in Table S2. Signal was detected with the Super Signal West Dura Extended Duration Substrate (Thermo Scientific, 34075) with a LI-COR Odyssey Fc imaging system (LI-COR). All figures are representative blots from at least three independent experiments unless otherwise stated.

### 2.7. Immunocytochemistry/FISH

Cells were cultured on 8-well polymer coverslips (Ibidi, 80826). For immunocytochemistry, cells were fixed with 4% PFA (H3K9Me3) or cold methanol (yH2A.x/nucleolus bright), permeabilized with 0.5% Triton X-100, and then blocked with 5% BSA. Slides were incubated in primary antibody at 4℃ overnight, and secondary at room temperature for at least 2 hours. Antibody information can be found in Table S2. FISH was performed with the HCR Gold RNA-FISH kit, probes, and X3 647 hairpins from Molecular Instruments according to the manufacturer’s protocol for fixing and staining cells in chambered cover slips. Cells were counterstained with Hoechst and Nucleolus Bright Red (Dojindo, N512) where appropriate for 10 minutes. All fluorescence images were captured at 60X on the Keyence BZ-X800. Imaging processing was performed evenly across all presented images, as follows: H3K9Me3 images were sharpened in Inkscape v.1.4.2; yH2A.X and nucleolus bright images were black balanced with Keyence; no additional processing was required for FISH (settings and raw images available upon request). All shown images are representative fields from at least three independent experiments unless otherwise stated. For quantification, images were imported to Image J, converted to RGB, color channels were split, and thresholding on the appropriate channel was performed to capture specific signal. Measurements were obtained for mean gray value and integrated density, limited to the set threshold.

### 2.8. EV and EV analyte isolation from cell culture supernatants

Cell culture supernatants were harvested on day 3 from proliferative control cells and day 7 from irradiated cells. Irradiated cells received a full media change on day 3. 10 mL supernatant was centrifuged and then concentrated to ∼500 µL by ultrafiltration with a 100K molecular weight cut off (Thermo Scientific, 88533). The resulting filtrate was separated with the Izon qEV original 70nm columns (Izon, ICO-70) and fractions 7-11 were collected (500 µL fractions). For downstream protein analysis, the isolated fractions were further concentrated by ultrafiltration (Thermo Scientific, 88524) for western blot or by speedvac for proteomics. For downstream RNA analysis, isolated fractions were processed through exoRNeasy (Qiagen, 77144), per manufacturer’s instructions.

### 2.9. ExoView

SEC fractions were analyzed with the ExoView R100 using an exosome human tetraspanin kit (Unchained Labs, 251-1044) and an automated platewasher. Samples were diluted 1:10 with incubation buffer, and chip processing and scanning were performed according to manufacturer directions. Data analysis was performed in the ExoView Analysis Software.

### 2.10. Patient plasma collection and EV RNA isolation

Patients were consented and samples were acquired as approved by the Baylor Scott & White Health institutional review board (020-325). Whole blood was collected in K2 EDTA tubes with subsequent (single-spin) plasma isolation. Plasma was stored at −80℃ until processing. EVs and RNA were directly extracted from 1mL patient plasma by exoRNeasy, per manufacturer’s instructions.

### 2.11. qRT-PCR

Whole cell RNA was isolated by RNeasy with manufacturer’s instructions (Qiagen, 74106) and EV RNA was isolated by exoRNeasy with manufacturer’s instructions (Qiagen, 77144). qRT-PCR was performed with the iTaq^TM^ Universal SYBR Green One-Step Kit (Bio-Rad, 1725151). Primer information can be found in Table S3. For SASP, analysis was performed with the 2^-ΔΔCt^ method (β-actin as housekeeping). For EVs, there is no well-accepted housekeeping RNA to perform normalization^40^. Therefore, relative expression was evaluated by calculating 2^-ΔCt^ (senEV-naiveEV) and weighting this expression by the ratio of input RNA (ng_naiveEV/ng_senEV), which was acquired by Qubit HS-RNA (Invitrogen, A32852) of the prepared RNA dilutions used during qRT-PCR. A synthetic cel-miR-39 cDNA (Norgen, 59000) was spiked in at a 1:10 dilution to identify any pipetting errors or presence of PCR inhibitors.

### 2.12. Total RNA library preparation and sequencing

Whole cell and EV RNA were cleaned with on-column DNase digestion (Zymo, R1013). RNA quality and quantity were determined with high-sensitivity RNA screentape (Agilent, 5067-5579). Total RNA libraries were prepared with SMART-Seq Total RNA Pico Input with ZapR Mammalian rRNA depletion and unique dual index kits (Takara, 634357 and 634752). Whole cell samples were prepared with 5 ng RNA, a 4 minute fragmentation, 5 adapter ligation cycles, and 14 PCR cycles. EV samples were prepared with 250-500 pg RNA, no fragmentation due to RNA size distributions, 5 adapter ligation cycles, and 18 PCR cycles. Library size was determined with high sensitivity D5000 screentape (Agilent, 5067) and quantity was determined by qPCR (Roche, KK4873). Equimolar pools with 5% PhiX were sequenced on the NovaSeq X Series platform. Paired-end sequencing was performed with 11 index cycles and 151 read cycles with NovaSeq Control Software 1.2.2.48004. Illumina BCL Convert v4.2.7 was used to convert base call (BCL) files into FASTQ files.

### 2.13. RNAseq analysis

High duplication rates are a known issue with low-input library prep^41^. However, deduplication without UMIs can cause increased noise and false negatives with little gain in accuracy^42–44^; this analytical risk is likely exacerbated in our non-fragmented libraries, as without random fragmentation, all identical biological transcripts with n >1 will appear indistinguishable from technical duplicates. Given these considerations, we proceeded without deduplication. Combined Read 1/Read 2 FASTQs were trimmed with cutadapt v4.4, and rRNA removed with sortmerna v4.3.4. Non-rRNA reads were quasimapped with Salmon v1.10.1 against a non-redundant genome annotation combined from GENCODE v40 and LNCipedia5.2, described originally by Rodosthenous *et al.*^45^. Transcript abundances were imported and converted to gene-level count matrices with tximport v1.18.0 in Rstudio 4.3.0. DESeq2 v1.42.0 provided normalized counts, and genes with less than 10 total reads across samples were removed. Differential expression analysis was performed with the Wald test with Benjamini-Hochberg correction. Designs included covariates of cell line/patient. Shrunken log_2_fold-change (log_2_FC) values were obtained with apeglm v1.12.0^46^. Additional analyses were performed in R v4.5.2. BiomaRt v2.62.1 was used to pull biotypes of the significant genes. Gene ontology over-representation was performed with clusterProfiler v4.14.6 enrichGO^47^. Gene set enrichment analysis (GSEA) was performed against normalized counts extracted from DESeq2 (averaged across biological replicates, where applicable) with the Broad GSEA 4.3.3 software using the Signal2Noise ranking and gene set permutation with H and C2 MSigDB collections.

### 2.14. Mass Spectrometry and Analysis

Protein from EV lysates [24µg (GBM43), 16µg (GBM102) and 1.6µg (A172)] were solubilized in a 2% sodium deoxycholate-based lysis followed by in-solution trypsin digestion as reported previously^48^. Data were acquired on RSLCnano U3000 LC system coupled to a Thermo Orbitrap Eclipse mass spectrometer with FAIMS Pro interface. 500ng peptides from each sample were directly loaded on a 50 cm * 75 µm ID C18 column (EasySpray ES903 columns) and separated by a 2 hr. LC-MS/MS method. Full scans were acquired in the Orbitrap at a resolution of 120,000 over a mass range of 375-1500 m/z. Using Data-Dependent Acquisition, most abundant precursors from the full scan were fragmented by HCD (Collision Energy of 32%) and detected in the ion-trap. FAIMS Compensation Voltage (CV) of −40/-60/-80 was enabled with 1 sec cycle time per CV. Raw spectra were queried in Proteome Discoverer (ThermoFisher Scientific, v2.4) using Mascot search engine (MatrixScience, v2.7) against a human protein database (UP000005640, downloaded Oct 2020) with the following parameters: tryptic peptide cleavage with up to two missed cleavages, oxidation of methionine as a variable modifications and carbamidomethyl on cysteine as a fixed modification, match between runs disabled. Peptide and peptide spectral matches (PSMs) were filtered using a false discovery rate of less than 1%, calculated using Percolator. The resulting protein abundances from the raw spectra search were normalized using variance stabilizing normalization in R (v.4.1.2). Significance was determined with Welch’s t-test followed by Benjamini-Hochberg (q-val < 0.05, |log_2_FC| > 2). GSEA was performed with Broad’s GSEA 4.2.3 software and then visualized using Cytoscape v3.10.2.

### 2.15. Graphing and Statistics

RNAseq and mass spectrometry analyses were performed as formerly indicated. All other graphing and statistical analyses were performed in GraphPad Prism v10.1.1, and results are shown as mean +/- the standard deviation of 3-5 independent experiments unless noted otherwise. Specific tests performed are indicated in the appropriate figure legends. Final figures were created with Inkscape v.1.4.2.

## 3. RESULTS

### 3.1. Radiation promotes senescence across GBM patient-derived models

To confirm that senescence is a relevant drug-induced functional state in GBM, we examined the response to radiation (IR) in 5 GBM cell lines (Table S1). We evaluated sensitivity to 2-10GY IR by relative viability at 24 hours, 72 hours, and 7 days (Figure 1A). Notably, effects were minimal until 7 days, suggesting a more delayed response like senescence than immediate cell death. This was supported by dose- and time-dependent effects on cell growth and senescence-associated-β-galactosidase (SA-β-gal) staining in the A172 cell line (Supplemental Figure 1A and 1B). Based on these data, we chose to further characterize senescence induction in all 5 GBM lines on day 7 following 6GY. As shown by live-cell imaging, 6GY did not result in large drops in cell viability but rather a sustained growth arrest through at least day 7, with most lines undergoing eventual proliferative recovery (Figure 1B); this is consistent with growth patterns documented in other models of tumor cell TIS^7^. Day 7 IR-treated cells were also larger and more granulated (Figure 1C), had increased SA-β-gal activity (Figures 1C and 1D), and showed increased H3K9Me3 expression, suggestive of SAHF (Figure 1E). Further, we observed downregulated PCNA (Figure 1F, top panel), upregulated p21 in a p53-dependent manner (Figure 1F, bottom panel), and upregulated IL6, IL1B, and MMP3, although this upregulation reached variable statistical significance (Figure 1G).

**Figure 1.**
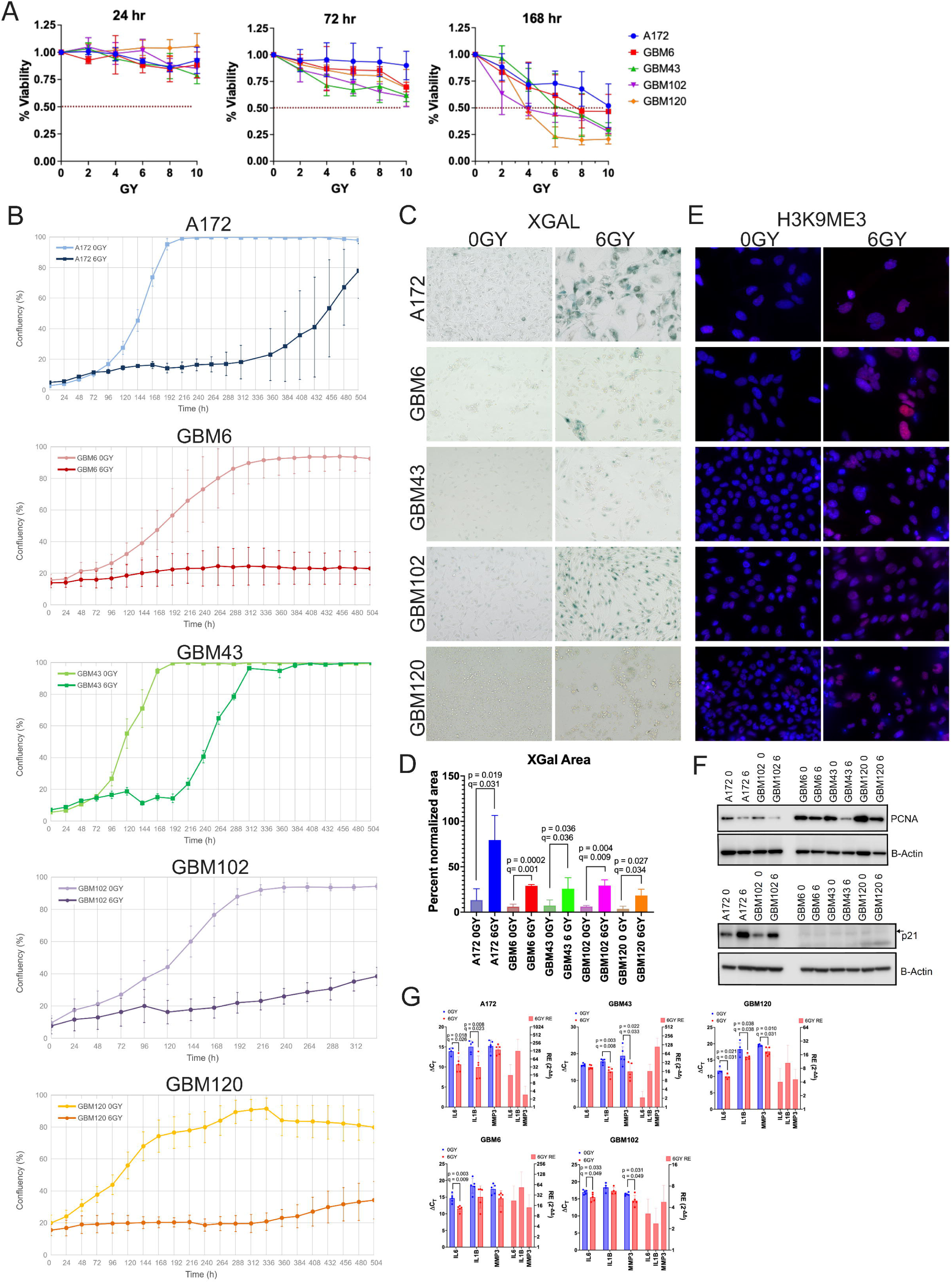
Evaluation of phenotypic markers of RIS following 6GY IR in GBM cells. **A)** Dose responses for five GBM cell lines following 0-10GY radiation at 24 hours, 72 hours, and 168 hours (D7) post-exposure, demonstrating that response to IR is delayed. The dotted lines mark 50% relative viability. **B)** Live-cell imaging demonstrating growth arrest in 6GY treated GBM. Graphs are shown as the mean confluency +/- SD of individual wells from a representative experiment (the chosen representative experiment for A172 is shown here and in supplemental figure 1). **C)** Brightfield microscopy shows phenotypic alterations in D7 treated GBM and increased SA-β-gal activity. **D)** Image J quantification of image area positive for XGal signal, normalized against cell area. Graph is mean +/- SD of at least 3 independent experiments. Significance was tested by unpaired t-test and Benjamini-Hochberg correction with FDR of 5%**E)** H3K9ME3 immunofluorescence reveals increased SAHF on D7. **F)** 6GY leads to reduced PCNA and cell-line/p53 dependent increases in p21. **G)** SASP factors are increased in all 5 GBM lines post-radiation. ΔCt bar graphs (left axis) are shown as mean +/- SD with independent experiments represented by data points. Relative expression (right axis) is shown as mean +/-SD across experiments. Significance was tested by unpaired t-test and Benjamini-Hochberg correction with FDR of 5%.

Finally, we profiled IR-induced transcriptional changes in A172, GBM43, and GBM102 by gene ontology and GSEA. Overrepresented gene ontologies in IR conditions (based on species with q-val <0.05 and log2FC>0), belonged to biological processes associated with viral response, interferon response, and cell size and organization (Figure 2A, left panel). These gene ontologies in response to radiation have been previously observed^49–51^. As expected for senescent cells, we found that underrepresented ontologies were associated with proliferation arrest, including DNA replication, chromosomal segregation, and cell cycle transitions (Figure 2A, right panel). GSEA further identified that post-IR cells were enriched in cell growth and cell stress hallmarks (Figure 2B, Table S4a), as well as multiple senescence-specific gene sets (Figure 2C, Table S4b). Altogether, our data supports robust RIS at both the transcriptomic and phenotypic levels, confirming the relevance of this phenotype in GBM and enabling biomarker discovery.

**Figure 2.**
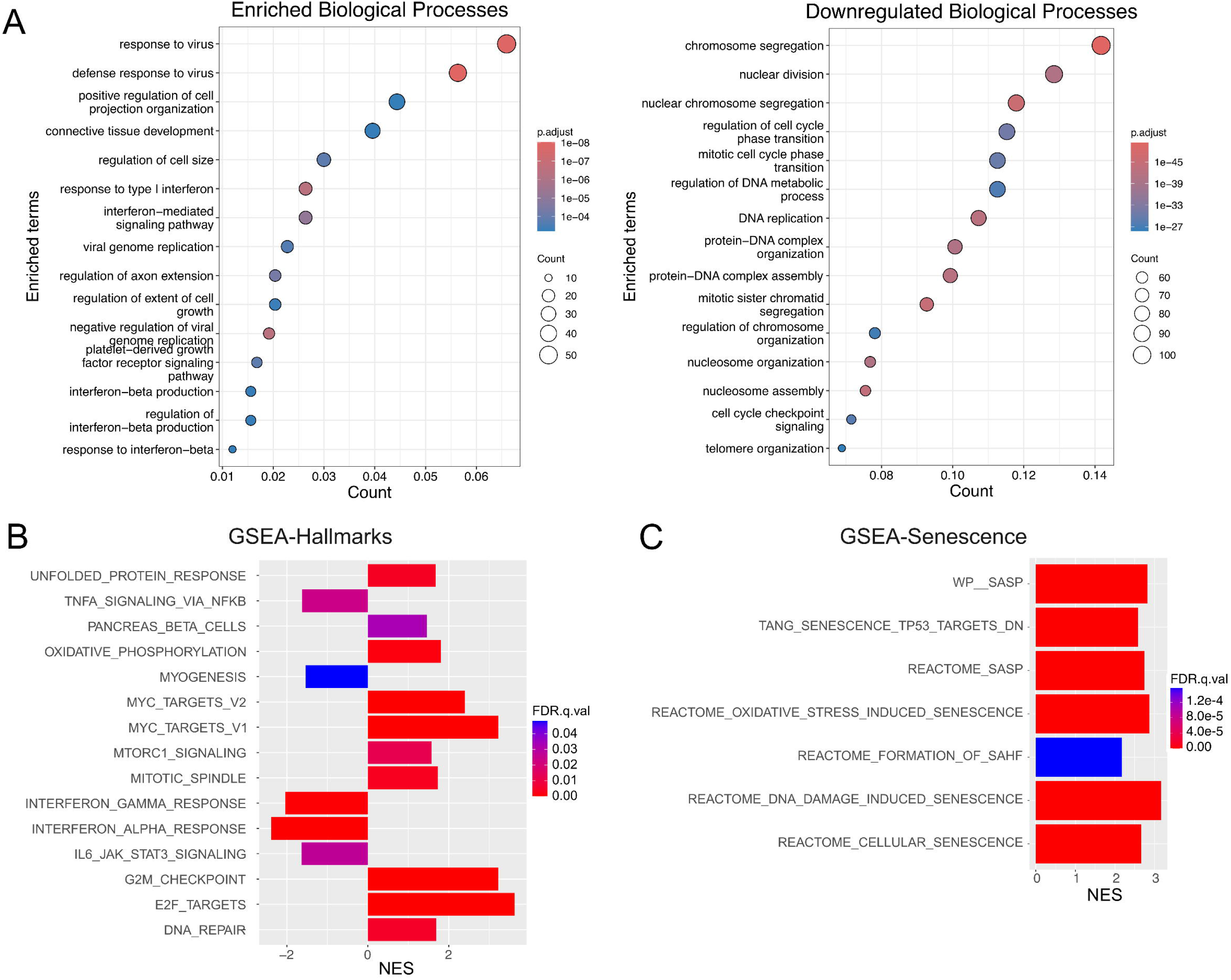
Profiling the senescent transcriptome in GBM. **A)** Significant differentially expressed genes (q-val <0.05) were queried for overrepresented (log2FC>0) and underrepresented (log2FC<0) gene ontologies with enrichGO. **B and C)** GSEA identifies expected gene sets for senescent cells. Significant hallmark gene sets are shown in **B,** and significant senescence-specific gene sets are shown in **C.** (NES = Normalized Enrichment Score)

### 3.2. GBM RIS is accompanied by elaboration of EVs that reflect the senescent phenotype

To identify potential signatures of RIS in GBM-derived EVs, we harvested EVs from treatment-naive, proliferative cells (naiveEVs) and senescent cells (senEVs) in our five GBM lines. EVs were isolated by size-exclusion chromatography prior to RNA analysis to ensure a high degree of purity and enrichment of desirable small exosome-like EVs (Supplemental Figures 2A-C). EV cargo includes a variety of RNA types^52^, and while long intact mRNAs have been identified, GBM EV mRNAs have previously been shown to be mostly fragmented^53^. This was also observed in our sample traces (Supplemental Figure 2D). As we were interested in both small noncoding RNA (<200nt) and mRNA markers of senEVs, we chose a total RNA approach with no fragmentation to minimize fragmentation-driven loss of small-to-mid length RNAs. While this approach does bias against intact long species, we posited that this would give us a broad view of putative senEV markers.

As expected, EVs clustered based on cell line of origin, with some separation between naiveEVs and senEVs for each line (Figure 3A). Still, when controlling for the effect of cell identity, we identified 580 differentially expressed species (q-val <0.1), though many of these had low magnitude changes (Figure 3B, Table S5). Of these, 477 (82.2%) belong to the Ensembl biotype “protein coding”. To identify whether the senEV transcriptome recapitulates that of senescent cells, we identified overrepresented gene ontologies in our significant species and enriched gene sets across the entire transcriptome. Overrepresented biological processes in the senEV group were related to protein assembly and processing, MHC processing, and cartilage/collagen (Figure 3C, left panel); there was minimal overlap between these and the top biological processes we identified in our whole cell transcriptomes (Figure 2A, left panel). However, downregulated DEGs belonged to biological processes associated with chromosomal and nucleosome organization, echoing what we found in the whole cell transcriptomes and what may be expected during senescence (Figure 3C, right panel). GSEA further revealed significantly enriched G2M checkpoint, E2F targets, and mitotic spindle gene sets in the senEV transcriptomes (Figure 3D, Table S6a). Of most interest, senEVs were also significantly enriched in three senescence gene sets (Figure 3E, Table S6b). We identified 23 species belonging to these senescent gene sets, as well as the broader Cell Age senescence database (build 2)^54^, that met both a q-value threshold of <0.1 and had log_2_FC ≥ 1 in either direction (Table 1). Of these, 17/23 changed in the expected direction based on the gene set(s) to which they belong. Overall, these combined data suggest that senEV transcriptomes are reflective of–though not identical to– the senescent cell population from which they were derived.

**Figure 3.**
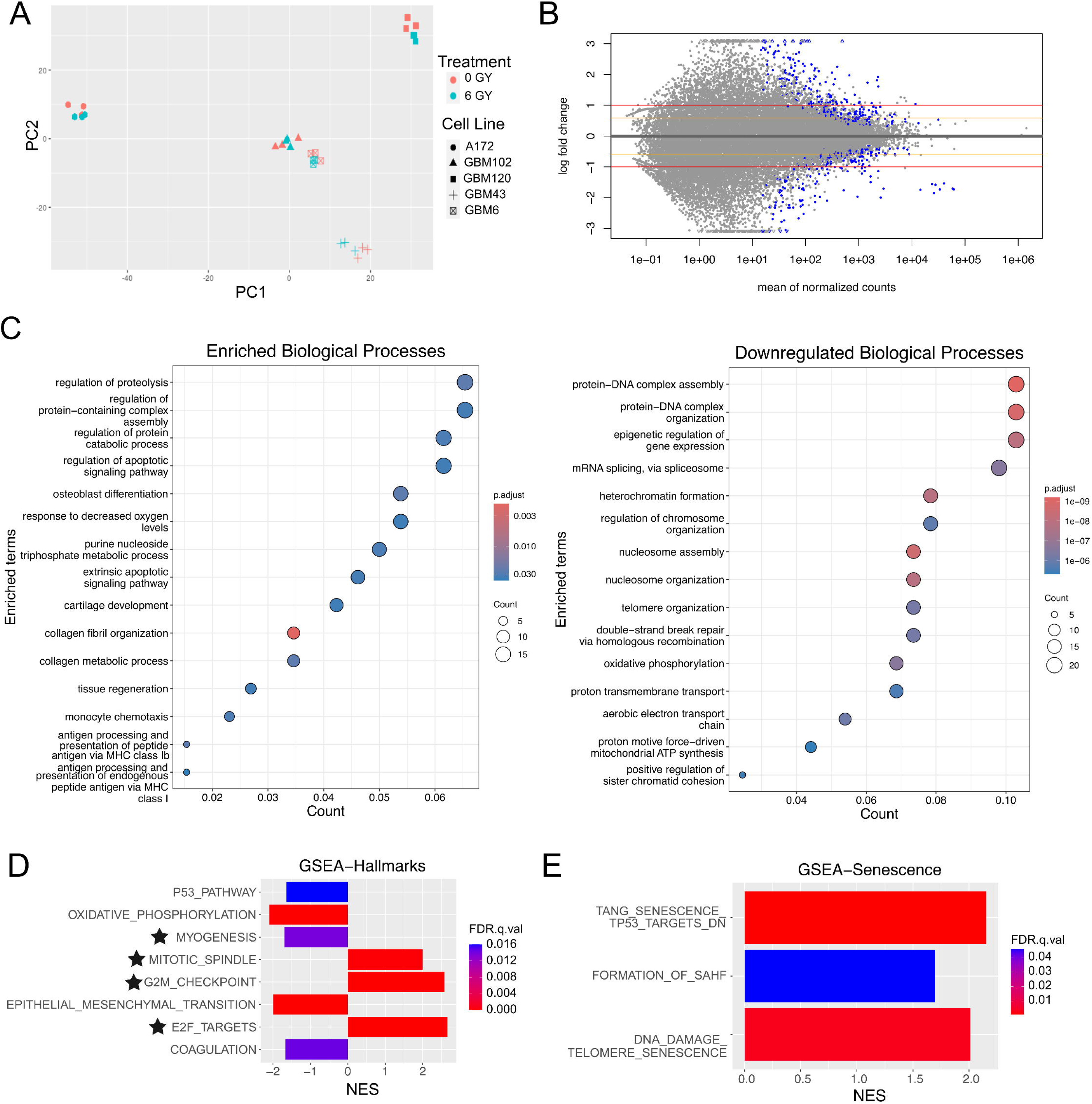
EVs secreted by RIS GBM cells have distinct transcriptomes from those secreted by treatment-naive GBM cells. **A)** Principal component analysis following variance-stabilization transformation reveals that variation by EVs is driven primarily by cell line of origin, though there is some distinction between senEVs and naiveEVs within cell line. **B)** MA plot, with significance indicated by blue color, showing that the majority of identified species have either low magnitude log2FC or do not reach significance. Red lines are drawn at Log2FC= −1,1 and orange are drawn at Log2FC= −0.585, 0.585. **C)** Queries of significant DEGs, as performed in Figure 2A, showing overrepresented and underrepresented gene ontologies. Note the similar underrepresented gene ontologies, right panel, as in the whole cell transcriptomes. **D and E)** GSEA identified hallmark gene sets that echo those of the whole cell transcriptomes during senescence, as well as significantly enriched senescence gene sets. Stars on D indicate gene sets found enriched with similar directionality in whole cell transcriptomes. (NES = Normalized Enrichment Score)

**Table 1.** Senescence associated DEGs in senEVs. DEGs were screened against the gene sets found to be enriched in senEVs, as well as the larger CellAge Expression Database. Species with opposing directionality of expected are shaded gray.

| Gene | Log2FC | shrunk<br>Log2FC | q-val | Gene Set |
| --- | --- | --- | --- | --- |
| IFI27 | 2.941 | 2.240 | 0.014 | CellAge_UP |
| SULF1 | 2.847 | 2.875 | 0.050 | CellAge_UP |
| CD163L1 | 2.653 | 1.435 | 0.060 | CellAge_UP |
| PPL | 2.326 | 1.742 | 0.014 | p53_correlated |
| BGN | 2.000 | 1.803 | 0.036 | CellAge_DN |
| APLP1 | 1.709 | 1.307 | 0.046 | CellAge_UP |
| COL5A3 | 1.680 | 1.376 | 0.016 | CellAge_DN |
| IGFBP5 | 1.609 | 1.441 | 0.000 | CellAge_UP, p53_correlated |
| NPDC1 | 1.496 | 1.362 | 0.000 | CellAge_UP |
| PRIM1 | -1.406 | -1.195 | 0.011 | CellAge_DN |
| CPPED1 | 1.362 | 1.046 | 0.060 | CellAge_UP |
| YEATS4 | -1.238 | -1.013 | 0.025 | CellAge_DN |
| H2AC4 | -1.142 | -0.875 | 0.044 | DNA_Damage_Sen |
| MKI67 | -1.139 | -0.972 | 0.012 | CellAge_DN |
| CDKN1A | 1.112 | 1.012 | 0.000 | CellAge_UP, DNA_Damage_Sen, SAHF |
| IGFBP3 | 1.074 | 0.749 | 0.078 | CellAge_UP |
| H2BC8 | -1.062 | -0.682 | 0.091 | DNA_Damage_Sen |
| LOXL2 | 1.061 | 0.692 | 0.084 | CellAge_UP |
| BAZ1A | -1.052 | -0.869 | 0.015 | CellAge_DN |
| H2AC16 | -1.049 | -0.884 | 0.011 | p53_correlated |
| CENPE | -1.019 | -0.808 | 0.025 | CellAge_DN, p53_correlated |
| H1-5 | -1.013 | -0.894 | 0.003 | DNA_Damage_Sen, SAHF |
| SMURF2 | -1.003 | -0.881 | 0.004 | CellAge_UP |

As an orthogonal approach to senEV biomarker discovery, A172, GBM43, and GBM102 EVs were subjected to mass spectrometry to evaluate protein cargo (Supplemental Figure 3, Table S7a-c). Interestingly, senEVs from A172 and GBM43 had increased numbers of unique proteins compared to naiveEVs (Supplemental Figure 3A). This increased protein abundance has been previously observed in EVs from senescent oral squamous carcinoma cells^55^. Previous work has also suggested that ATP6V0D1 and RTN4 may serve as biomarkers from senescent fibroblasts^37^; however, this was not found to be the case in our GBM senEVs, as we observed no appreciable upregulation of ATP6V0D1 and upregulation of RTN4 in only GBM43 (Supplemental Figure 3B). In fact, we identified no shared upregulated and only three shared downregulated proteins (q-val <0.05) across lines: CEP55, KIF23, and RACGAP1 (Supplemental Figure 3C). Additionally, we found only modest signals of senescence at the protein level in our senEVs, both when considering gene set enrichment (Supplemental Figure 3D) and individual significant species (p-val <0.05, none of which met q-val <0.05) (Supplemental Figure 3E). Given this, we concluded that the senEV transcriptome was a stronger analyte for senEV biomarker development.

### 3.3. A panel of snoRNAs are candidate species for senEV biomarker development

Encouraged by the signatures of senescence-associated RNA cargo in senEVs, we wanted to identify additional markers that strongly differentiate senEVs from naiveEVs. We ranked significant species by their shrunken |log_2_FC| values and examined the top 20 (Figure 4A, Table S8). Four species were protein coding, and two of these four were senescence-associated (*SULF1, IFI27*). Strikingly, of the 16 remaining non-coding species, 12 belonged to the snoRNA class. Of particular interest, *SNORA13* has recently been implicated in senescence^56^.

**Figure 4.**
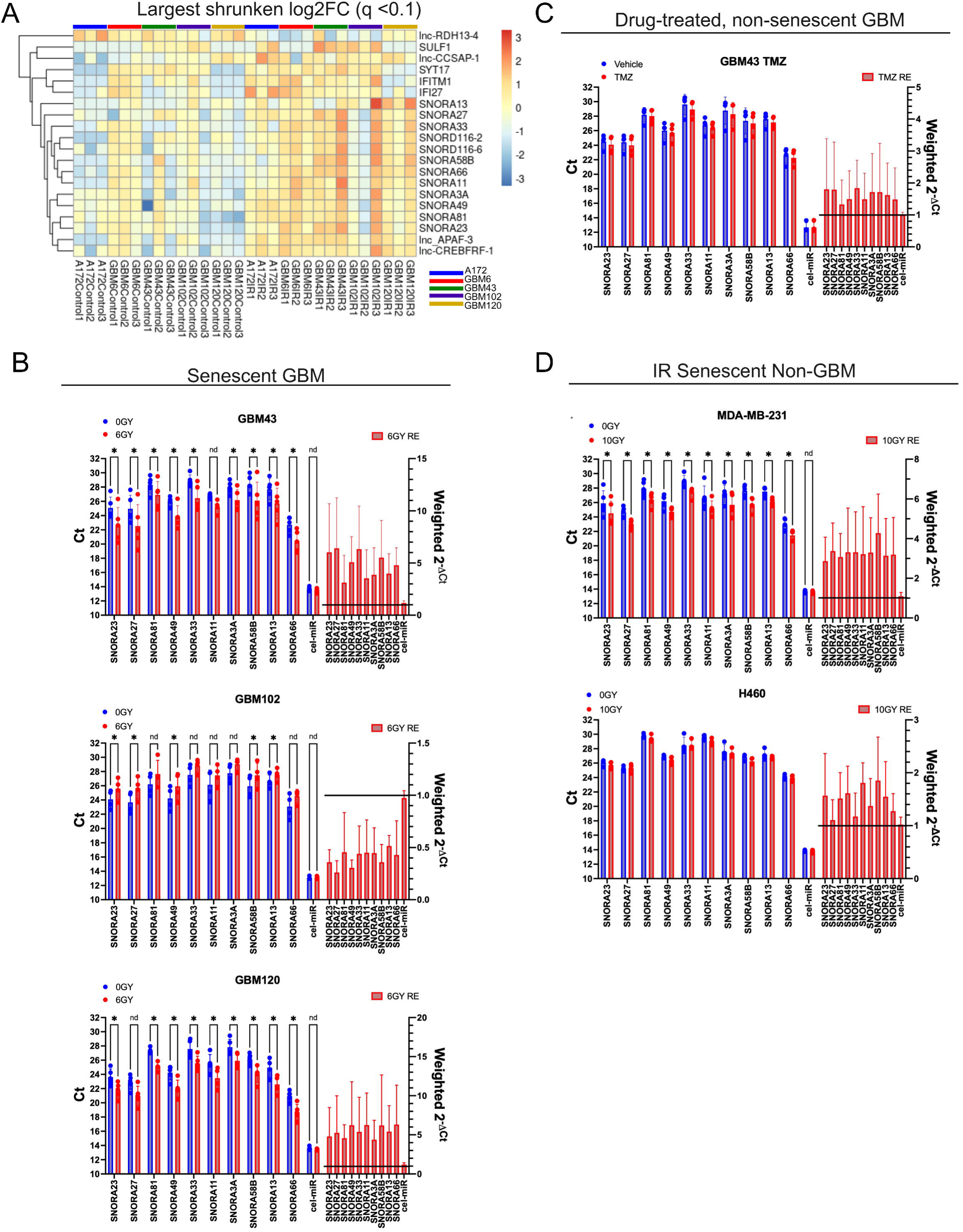
senEVs are generally enriched in snoRNAs. **A)** Heatmap of log2(normalized_count+1), scaled by row, for the top 20 species after ranking DEGs by shrunken log2FC values. **B)** snoRNA expression patterns observed in RNAseq were validated by qRT-PCR in GBM43, GBM102, and GBM120. cel-miR-39 cDNA was used as a qPCR spike in control. **C)** snoRNA were not significantly upregulated in non-senescent, TMZ treated GBM43 derived EVs. **D)** snoRNA abundance was evaluated in the senEVs from breast (MDA-MB-231) and lung (H460) cancer cell lines following RIS. For B-D, Ct values (left axis) and modified relative expression (right axis, see methods for details) are shown as mean +/- SD with independent experiments represented by data points. Significance was tested by paired t-test with Benjamini-Hochberg correction with FDR of 5% to account for variable input per replicate. * indicates q-value <0.05, n.d. indicates no discovery (q-value > 0.05).

To test the rigor of the RNAseq results, we performed qRT-PCR validation in an independent collection of GBM43, GBM102, and GBM120 EVs. Limited RNA yield from GBM6 and A172 EVs precluded evaluation, and primers for SNORD116-2 and −6 could not be built with confidence, so these were omitted in follow-up. As shown in Figure 4A, snoRNAs by RNA sequencing were upregulated in GBM43 and GBM120 senEVs, but not GBM102 (with the exception of one replicate). This was confirmed by qRT-PCR, as increased snoRNA expression was found in only GBM43 and GBM120 senEVs (Figure 4B). We also mined our proteomic data for snoRNA associating proteins (SRPs)^57^, which would suggest co-packaging and provide additional indirect evidence of snoRNA incorporation in senEVs. Indeed, we found SRPs enriched in, and unique to, GBM43 and A172 senEVs (Table 2), but not in GBM102 senEVs. Overall, these validated results argue for snoRNA enrichment as a putative biomarker for senEVs. While not shared by the GBM102 model, we consider the enrichment of snoRNA across ⅘ models is likely appropriate for capturing a molecularly diverse functional state in a very heterogenous tumor type^58–60^.

**Table 2.** snoRNPs in GBM EVs identified by mass spectrometry. Log2FC values represent change of senEVs compared to naiveEVs. Shaded Log2FC boxes indicate that the protein was not identified in either condition, whereas shaded p-val and q-val boxes indicate that the protein was not identified in at least 2/3 of either condition, limiting significance testing.

|  | A172 |  |  | GBM43 |  |  | GBM102 |  |  |
| --- | --- | --- | --- | --- | --- | --- | --- | --- | --- |
| Gene | Log2FC | p-val | q-val | Log2FC | p-val | q-val | Log2FC | p-val | q-val |
| FBL | 1.138 | 0.216 | 0.325 | UNIQUE 6GY |  |  | UNIQUE 0GY |  |  |
| NHP2 | -0.391 | 0.053 | 0.123 | 2.836 |  |  |  |  |  |
| NOP56 | UNIQUE 6GY |  |  | 3.135 | 0.002 | 0.007 |  |  |  |
| NOP58 | UNIQUE 6GY |  |  | 3.204 | 0.007 | 0.018 |  |  |  |
| GAR1 | UNIQUE 6GY |  |  | 2.936 |  |  | UNIQUE 0GY |  |  |
| SNU13 | UNIQUE 6GY |  |  | 4.929 |  |  | UNIQUE 0GY |  |  |
| DKC1 | UNIQUE 6GY |  |  | UNIQUE 6GY |  |  |  |  |  |
| NOP10 |  |  |  | UNIQUE 6GY |  |  |  |  |  |

To further define the association of EV-snoRNA with the RIS phenotype, we tested whether snoRNA might increase under conditions of cell stress irrespective of senescence. In GBM43 cells, TMZ produces a response that is associated with minimal upregulation of SA-β-gal, suggesting limited senescence induction (Supplemental Figures 4A-C). Reflective of this limited senescence, upregulation of the snoRNA panel in EVs following treatment was insignificant (Figure 4C). To determine whether snoRNA upregulation might be observed in other tumor types, we quantified snoRNA enrichment in the EVs secreted by MDA-MB-231 and H460 cells induced to senescence by 10GY IR (Supplemental Figure 4C). While we only observed a moderate upregulation in H460 senEVs, all snoRNAs were significantly upregulated in MDA-MB-231 senEVs (Figure 4D). This data further supports this panel of snoRNAs as putative markers for senEVs and expands their potential utility beyond GBM.

### 3.4. GBM patient plasma EVs post SOC support further development of a senEV bioassay

Translation of a biomarker for clinical RIS detection in GBM will be difficult without commensurate IHC approaches to assess senescence in post-treatment biopsied tissue. However, signals of RIS-associated gene sets and/or snoRNAs in patient plasma EVs after completion of radiotherapy encourage future studies and motivate development of senEVs as a biomarker for RIS in patients. Therefore, we performed total RNA sequencing on EVs from 4 paired GBM patient plasma samples collected at time of surgical resection (PreOP) and following completion of 46GY IR with concurrent temozolomide (PostSOC). Patient demographics are included in Table S9.

Interestingly, plasma EV transcriptome profiles showed greater dispersion between PreOP samples and less dispersion between PostSOC samples (Figure 5A). While this is potentially aligned with prior work showing that patient PBMCs exposed to radiation *ex vivo* cluster by dosage and not donor^61^, we further examined the effects of final library size and sample storage duration as potential confounders. However, we did not find strong evidence supporting these factors as covariates (Supplemental Figure 5A and 5B). This clustering suggests that patient exposure to radiation and systemic temozolomide is associated with drastic cargo changes in plasma EVs.

**Figure 5.**
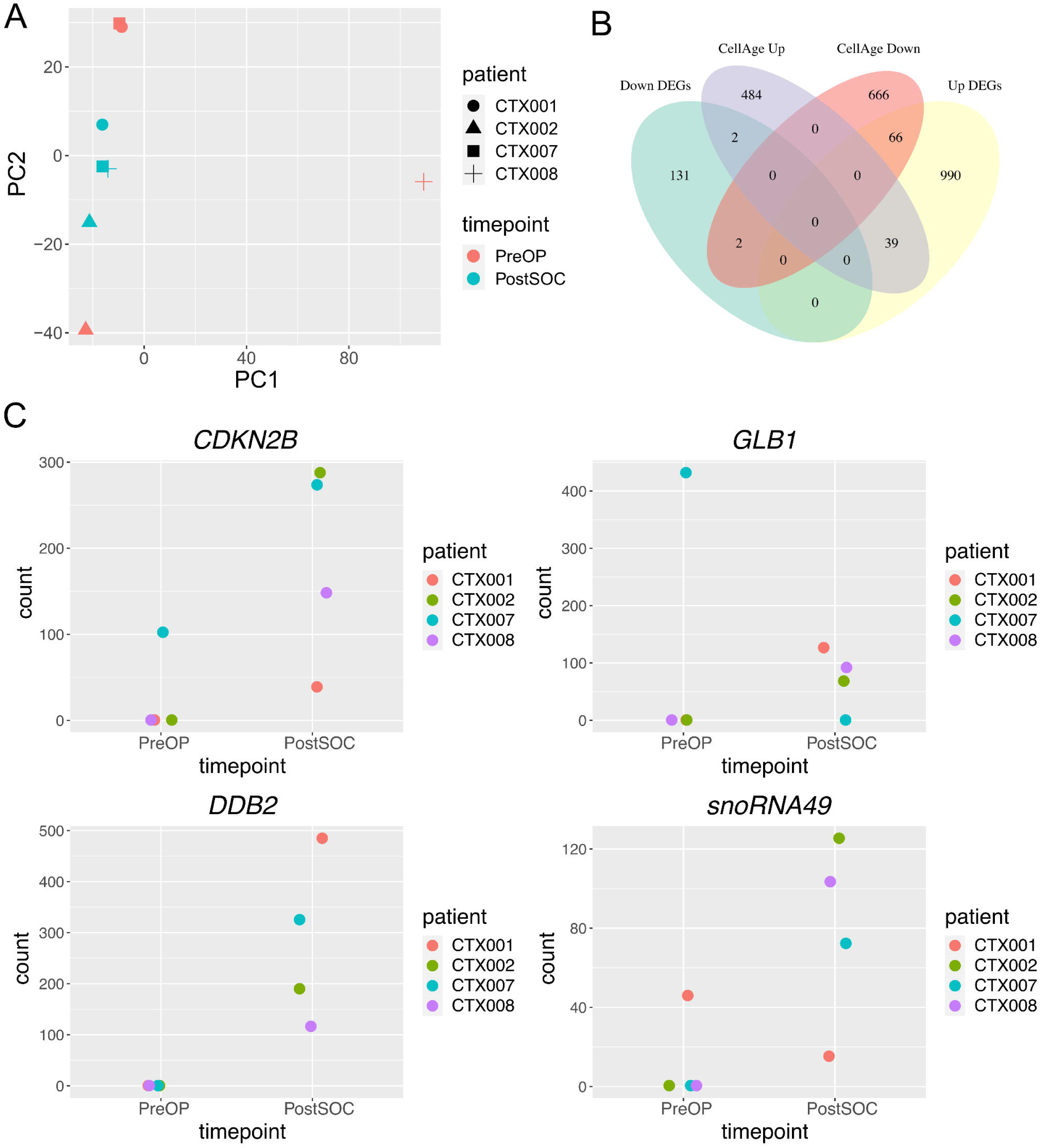
Species of interest are upregulated in GBM patient plasma EVs post-radiation. **A)** Principal component analysis following variance-stabilization transformation indicates clustering driven by treatment, particularly in postSOC EVs. **B)** Significant DEGs (PostSOC vs. PreOP) were screened against the Cell Age senescence expression database. 39 genes were observed to be upregulated as expected and 2 genes were observed to be downregulated as expected. **C)** Normalized counts for *CDKN2B, GLB1, DDB2,* and *SNORA49*.

We found that upregulated species in PosSOC samples with q-val <0.1 were enriched in biological processes associated with the immune system, particularly T-cell related, and cell cycle regulation (Supplemental Figure 5C). In fact, *CDKN1B*, which encodes for the cyclin dependent kinase inhibitor p27, was amongst the top most differentially expressed species (Supplemental Figure 5D). To determine whether elevated signals of senescence are present in the PostSOC EVs as a group, we again turned to GSEA. However, senescence gene sets were not significantly enriched (data not shown). We screened significant DEGs (q-val <0.1 and |log_2_FC| ≥ 1) against the CellAge database, theorizing that individual senescence-associated species may still be detectable, despite lack of whole gene set enrichment. 109 DEGs overlapped with the Cell Age Database, but a large percentage were discordant in their directionality (Figure 5B). We identified 41 species that aligned with CellAge (Figure 5B and Table S10). This includes well-known *CDKN2B* (the cell cycle inhibitor p15), for which we observed increased abundance in 4/4 paired samples; *GLB1* (β-galactosidase), for which we observed increased abundance in 3/4 paired samples; and *DDB2* (damage specific dna binding protein 2), for which we observed increased abundance in 4/4 paired samples (Figure 5C). We acknowledge the absence of counts for these genes in some samples limits confidence in the extent of upregulation, which is reflected in their low shrunken log2FC values (Table S10); however, given that the genes are not lowly expressed where detected (normalized counts > 10), we believe that the trend towards upregulation is likely real.

Finally, we queried DEGs for snoRNAs and identified three species: *SNORA49, SNORA22, SNORA77A*, of which the two former were upregulated. Notably, *SNORA49* was identified in our *in vitro* senEVs, as well. 3/4 patients showed increased *SNORA49* normalized counts in their PostSOC sample compared to their PreOP (Figure 5C). Of interest, the patient that did not show upregulated *SNORA49* is the only patient with definitive radiographic evidence of progressive disease (Table S9). While these data are not evidence of a snoRNA or senescence enrichment, per se, we find that the increase in detectable species of interest in PostSOC EVs from an unenriched plasma source with an untargeted sequencing effort is encouraging.

### 3.5. snoRNA enrichment in senEVs is independent of drastic nucleolar alterations

Prior work in the MCF7 breast cancer line has shown that radiation-induced senescence results in nucleolus fragmentation^62^. As such, we questioned whether this disruption to the nucleoli might cause wide-scale release of snoRNA into the cytoplasm, promoting increased capture and packaging into EVs. We examined the nucleoli in the four GBM cell lines exhibiting upregulated senEV snoRNA abundance by Nucleolus Bright Red, which stains RNA with a preference for rRNA in the nucleolar regions; nucleolar staining presents as distinct large “puncta” like regions within the nucleus while non-rRNA staining presents as more unrefined cytoplasmic/perinuclear stain. However, we observed no nucleolar fragmentation (Figure 6A), even in individual nuclei displaying persistent DNA damage/ sustained DNA damage response signaling (Figure 6B, example nuclei indicated with arrows throughout). Accordingly, we found no mass release of *SNORA49* or *SNORA13* out of the nucleolar regions when visualized by fluorescent *in situ* hybridization (FISH) (Figure 6C and Supplemental Figure 6). Although there appeared to be a trend in both average pixel intensity (mean gray value) and total abundance (integrated density) of *SNORA49*, these increases generally did not reach statistical significance, likely due to large variance between biological replicates (Figure 6D). This is consistent with results from qRT-PCR in whole cell lysates across a subset of the snoRNA panel (Figure 6E). Altogether, this data suggests that senEV snoRNA enrichment is not a function of nucleolar region collapse, but may perhaps be instead related to a closely controlled equilibrium of cellular abundance during senescence.

**Figure 6.**
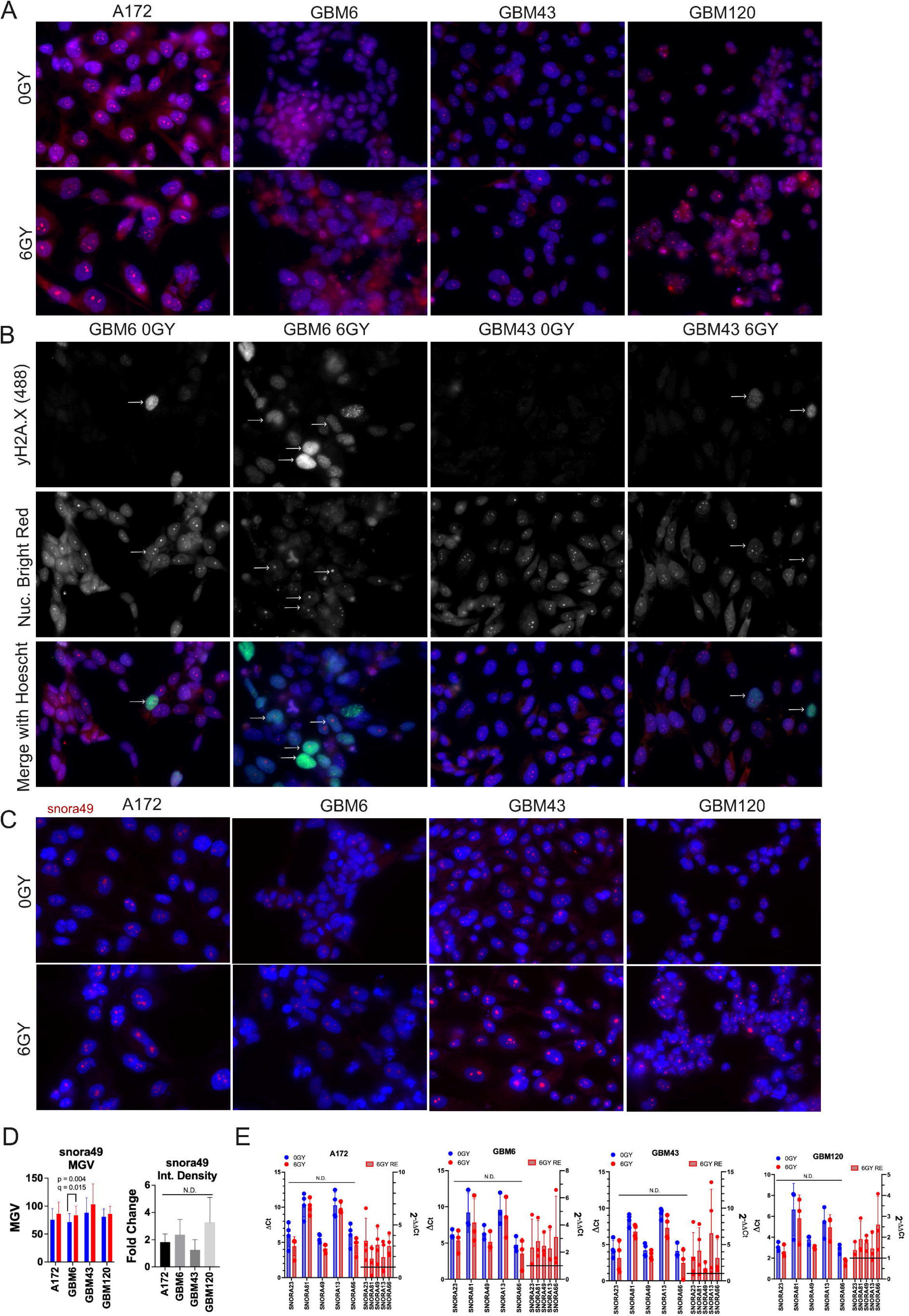
Examination of nucleoli and snoRNA localization during senescence. **A)** Nucleolus bright red staining shows no appreciable nucleolar fragmentation (increased number of small, misshaped nucleoli within the nucleus) in senescent GBM cells. **B)** Co-staining of nucleolus bright and yH2A.X reveals that nucleoli remain intact in cells with persistent DNA damage. White arrows mark nuclei of interest across the presented channels and merged images **C)** *SNORA49* is still abundant in and primarily localized to the nucleolar regions during senescence. **D)** Quantification of pixel intensity (mean gray value, left) and abundance (integrated density, right) for *SNORA49*. Graphs show mean +/- SD for three independent experiments. Significance was determined by ratio paired t-tests (on raw data values) with Benjamini-Hochberg correction, using FDR of 5%. **E)** qRT-PCR for a subset of snoRNAs in whole cell lysates. Significance was determined by unpaired t-test with Benjamini-Hochberg correction, using FDR of 5%.

## 4. Discussion

Radiation remains a standard of care for many solid tumor types, garnering temporal control but eventuating in recurrent disease. Interrogation of and interference with radiation-induced senescence (RIS) may offer a therapeutic opportunity to minimize residual disease following treatment. However, RIS– and therapy-induced senescence, in general– is temporally and spatially dynamic, meaning rigorous evaluation requires measurement across time points and regions. Current methodologies for evaluating senescence in patients are histochemical or transcriptomic, thereby requiring tissue^17,27,63,64^; for certain tumor types like GBM, this requirement precludes evaluation after SOC. The development of a minimally invasive liquid bioassay for RIS would open avenues for both an improved understanding of senescence clinically and the potential for a precision medicine approach for senotherapeutics. EVs have shown promise as biomarkers for binary findings like the presence or absence of GBM^29,65^, but we questioned whether EVs may also be able to capture more complex phenotype information. Further, while previous studies have examined cargo in EVs released by senescent cells, the focus has primarily been on functional biological outcomes of EV release and uptake, rather than identifying to what extent EV cargo reflects or can report on the senescent cell phenotype^33,66^.

We found here that EVs elaborated by senescent GBM contain transcriptomic features that may enable their use as an analyte for RIS detection. Specifically, senEVs contain increased abundance of senescence-associated RNA species and decreased abundance of species that simultaneously are downregulated at the whole cell level. Unexpectedly, we also found that senEVs contain an enrichment of snoRNAs, which are likely packaged with their protein binding partners. The roles of snoRNA beyond rRNA modification are understudied, but at least one snoRNA identified in our senEVs, *SNORA13,* has been implicated in senescence^56^.

It is important to emphasize that at least two separate classes of RNAs may identify senEVs (coding RNAs of senescence genes and the noncoding snoRNAs). Senescent cells are highly heterogeneous at the transcriptional level and phenotype development can be variable between cell lines and between senescence-stimuli^58,67,68^; as a result, identification of senescent cells generally requires assessment beyond any singular marker. A similar finding is made here in senEVs, as 2/7 senescent models failed to demonstrate upregulated EV snoRNA. While initial presentation in ∼70% of *in vitro* RIS models is promising, this suggests the need for refinement. This may be accomplished by either further stratification of tumor genotypes that associate with snoRNA upregulation, or by incorporating additional markers into the signature– i.e. the coding senescence RNAs– to expand capacity to capture this heterogenous cell state. We posit that the latter option, a combination of both senEV RNA classes, will offer the greatest sensitivity and specificity to detect senescence. Future assay development will likely incorporate a targeted sequencing panel against multiple of our identified RNAs, combined with an algorithmic or machine learning–based approach to establish robust thresholds for defining a ‘positive’ signal of senescence.

Excitingly, from four sets of longitudinal patient plasma samples we were also able to detect increased abundance of both senescence-associated and snoRNA species in post-SOC EVs compared to pre-operation EVs. While broad conclusions are limited due to minimal cohort size and lack of UMIs to enable more confident transcript quantitation, these initial results demonstrate the feasibility of detecting putative markers of cell stress and functional states from plasma EVs. Further, our approach here was performed without enrichment for brain or GBM-derived EVs. Given the systemic administration of temozolomide, it is likely that a portion of the detected signals reflect stress imposed on host blood and non-brain tissues. We believe these signals are still of importance to patient health, as senescence outside of the tumor tissue of origin has previously been implicated in therapy-associated side effects, such as chemotherapy-induced neuropathy and cardiotoxicity^69,70^. Rather than moving forward with enrichment of GBM/brain-derived EVs in future work, we posit that retaining signals of senescence in both GBM/brain and other host tissues will be beneficial. To this end, recent advancements in single EV sequencing^71,72^ may allow for the simultaneous detection of both senescence- and tissue-associated signatures, enabling a more delineating assessment of host v. tumor responses.

Perhaps the most difficult aspect of translating a senEV bioassay for RIS– or any clinical evaluation of RIS– will be the capacity to assess specificity and sensitivity of the markers. Future studies should certainly include larger patient cohorts with more robust time point acquisition, alongside matched radiographic response data. Tumor types that allow for repeated tissue biopsy, such as breast cancer, may therefore be an ideal additional patient cohort. This would enable concurrent standard detection of tissue senescence to allow for true measure of senEV capabilities. While preliminary, our results in GBM patients presented here support such an undertaking.

Finally, it is important to acknowledge that radiation promotes multiple functional states in GBM, including but not limited to autophagy, persister states that depend on lipid peroxidase pathways, and stem cell-like transitions^73–75^. Notably, these functional states are not mutually exclusive, and indeed, autophagy and senescence are closely intertwined^76^. While our current work has focused on RIS, the senEV signature likely represents only one opportunity to identify appropriate adjuvant therapies for GBM patients. Our data here provides proof-of-concept that EVs recapitulate and relay transcriptional information about complex processes that the originating cell population is experiencing. We posit that similar signatures can be developed for additional functional states, which would broaden the applications for a post-SOC EV assay for GBM and other cancer patients. While disentangling the markers for concurrent phenotypes will present a challenge, the data here ultimately supports extracellular vesicles as promising analytes for the development of liquid biopsies detecting drug-induced functional states, particularly therapy induced senescence.

## Supporting information

High Resolution Images

## 5. Acknowledgments

VD is supported by the American Cancer Society – Simone Charitable Trust - Postdoctoral Fellowship, PF-24-1198074-01-CDP. This work was supported in part by the Dorrance Research Fund, the Lori Lane & Andy Spryow Fund, and BSWH CP18. This research includes work performed in the TGen Collaborative Sequencing Center, a City of Hope Comprehensive Cancer Center supported shared resource (NCI-P30CA033572), and by the Integrated Mass Spectrometry Core at City of Hope Comprehensive Cancer Center, funded by the National Cancer Institute of the National Institutes of Health under award number P30CA33572. The authors thank critical members of the Mass Spectrometry Core, including Ritin Sharma, Brooke Lovell, and Melissa Martinez.

## 6. Data Accessibility

Accession numbers will be provided for RNA sequencing and mass spectrometry upon publication. Full western blot images are shown in Supplemental Figure 7. Raw microscopy images are available upon request.

## 8. Figure Legends

**Supplemental Figure 1. Additional conditions of RIS in GBM. A)** Live cell imaging and **B)** XGal staining across different doses and times in the A172 cell line.

**Supplemental Figure 2. Extracellular vesicle quality control. A)** Western blot against the endoplasmic reticulum protein calnexin and the exosomal tetraspanin protein CD9 across 3 independent EV lysates and one whole cell lysate from the GBM43 cell line. **B)** Particle counts for each capture spot on the exoview chip for GBM43 and GBM120 EVs following SEC. **C)** Size analysis of the captured particles by spot. N=1 independent experiment, but >1K particles per capture spot, per condition. **D)** Representative high sensitivity RNA tapestation traces showing the heavy fragmentation profile of EV RNA compared to whole cell RNA. While RINe scores are not always appropriate metrics for EV RNA, the abundance of fragmented RNA is clear.

**Supplemental Figure 3. Mass spectrometry of senEVs. A)** Number of proteins and peptides identified in A172, GBM43, and GBM102 naiveEVs and senEVs. Notably, in GBM43 and A172 senEVs there is increased protein content, as has previously been observed. **B)** RTN4 and ATP6V0D1, which were previously suggested as markers of senEVs in fibroblasts, were not observed to be significantly or consistently enriched in GBM senEVs. **C)** Venn diagrams showing significant (p<0.05) proteins identified in senEVs. **D)** GSEA identified enriched senescence gene sets in at least 1/3 lines, which while promising, appeared less convincing than the transcriptomic signals identified throughout the study. **E)** Venn Diagrams (top panels) of significant (p<0.05) proteins identified in senEVs belonging to the CellAge Database, and corresponding bar plots for proteins conserved across lines (bottom panels).

**Supplemental Figure 4. Additional conditions for snoRNA enrichment. A)** Dose response curve to temozolomide, read at 144 hours, in GBM43. Please note that this figure is shared between this manuscript and Mankame *et al* ^77^ as it was produced for dual use by co-author N. Tang. **B)** Live cell imaging for GBM43 cells with different doses of temozolomide (n=1). **C)** XGal images for the indicated conditions. Representative figures of 3 independent experiments.

**Supplemental Figure 5. Evaluation of postSOC versus preOP plasma EVs from GBM patients. A)** Separation/clustering of plasma samples did not appear to be heavily driven by final library size (top panel) or sample age (bottom panel). **B)** MA-plots with (bottom panel) and without (top panel) Apeglm shrinkage do not suggest improper normalization of library size. Significance is indicated by blue color, red lines are drawn at Log2FC= −1,1 and orange lines are drawn at Log2FC= −0.585, 0.585. **C)** Overrepresented biological processes in post-SOC plasma samples were primarily related to T-cell and replication associated ontologies. **D)** Heatmap of the top 20 differentially regulated genes with q-val <0.1. Of interest is the increased abundance in *CDKN1B*.

**Supplemental Figure 6. Snora13 *in situ hybridization.*** *SNORA13* localization during control (0GY) and senescent (6GY) conditions. A no probe control is shown to evaluate non-specific background. Representative images from 2 independent experiments. No quantification was performed due to n<3 replicates.

**Supplemental Figure 7. Uncropped western blots.**

## Notes

### Competing Interest Statement

The authors have declared no competing interest.

### Summary of Updates

Figure 1 has been corrected to include a panel that was originally missing. Additional new findings have been included. Some text has been updated as well. Major findings and conclusions remain the same between versions.

