## Supplementary figures and images for "Molecular profiling of glioblastoma-derived extracellular vesicles identifies small nucleolar RNAs as candidate liquid biomarkers for radiation- induced senescence"

### Figure 1.pdf

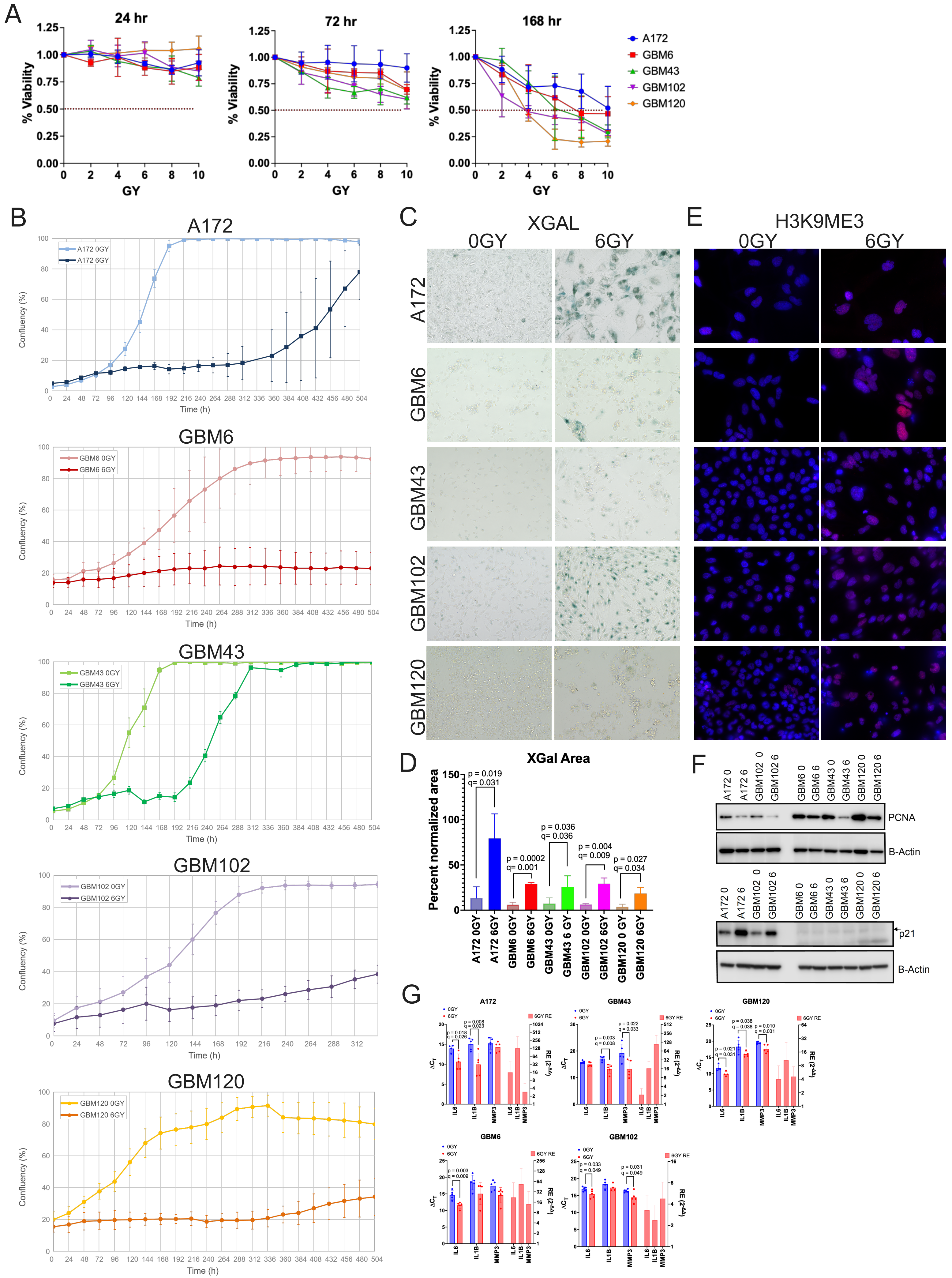

### Figure 2.pdf

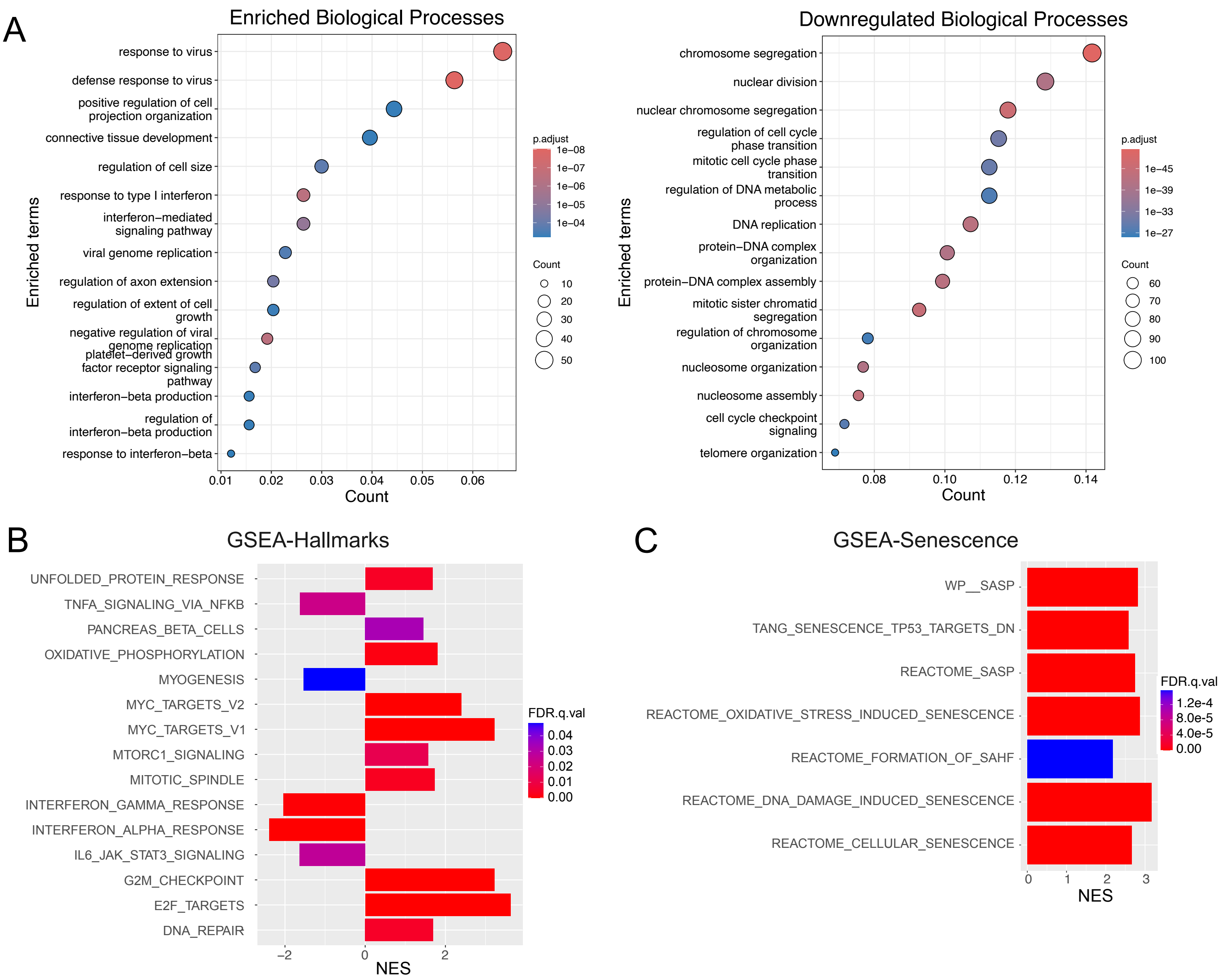

### Figure 3.pdf

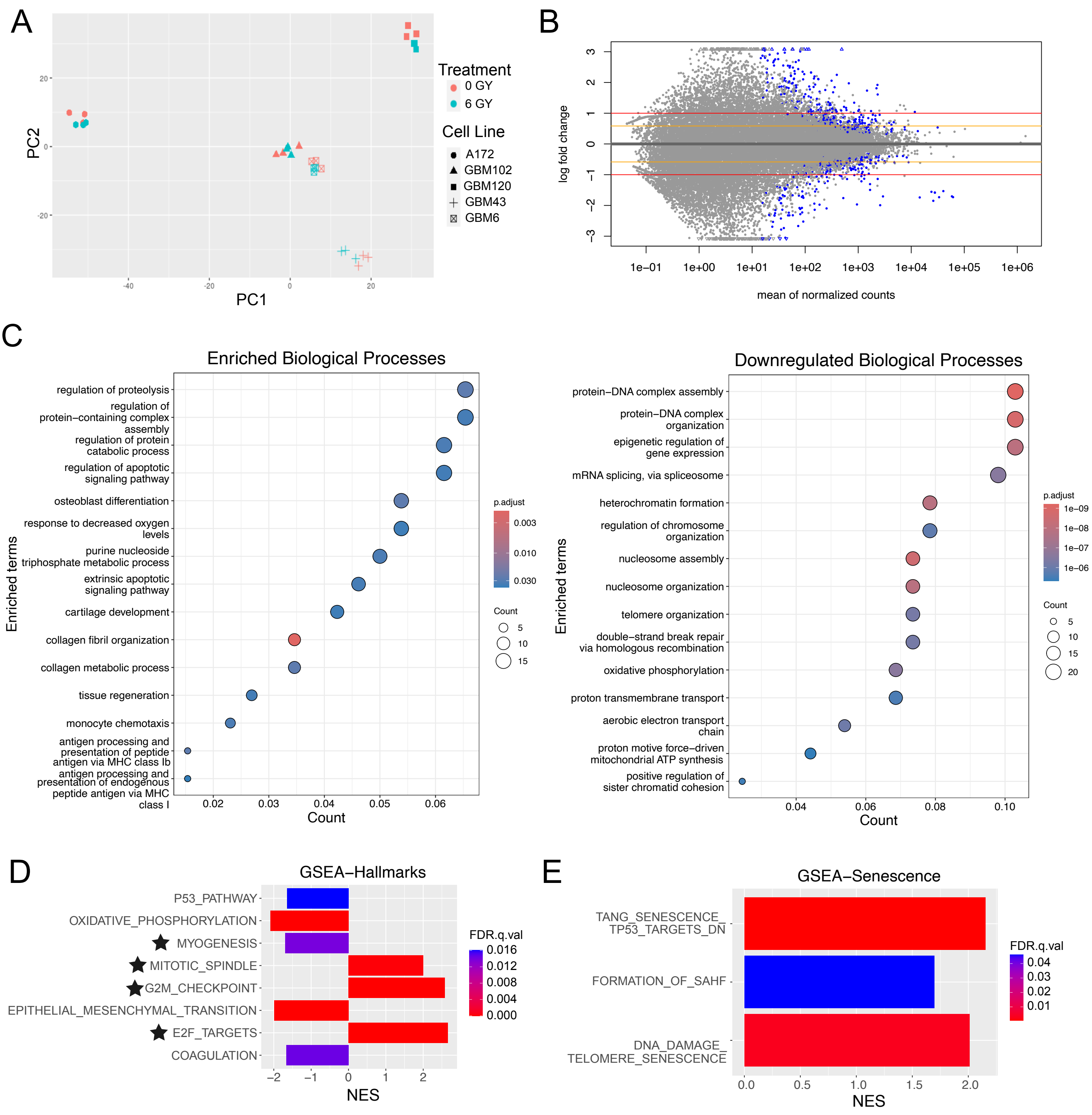

### Figure 4.pdf

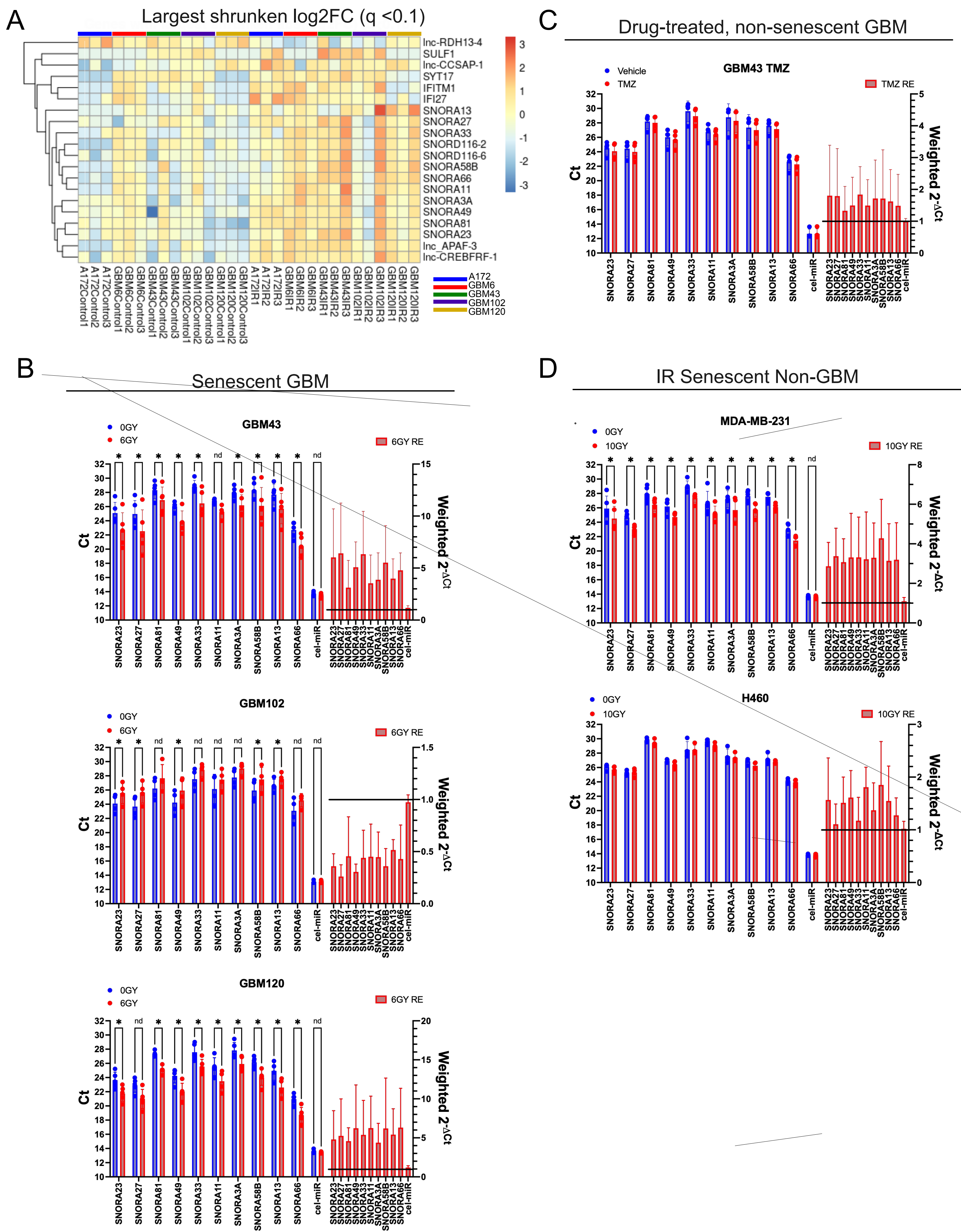

### figure 5.pdf

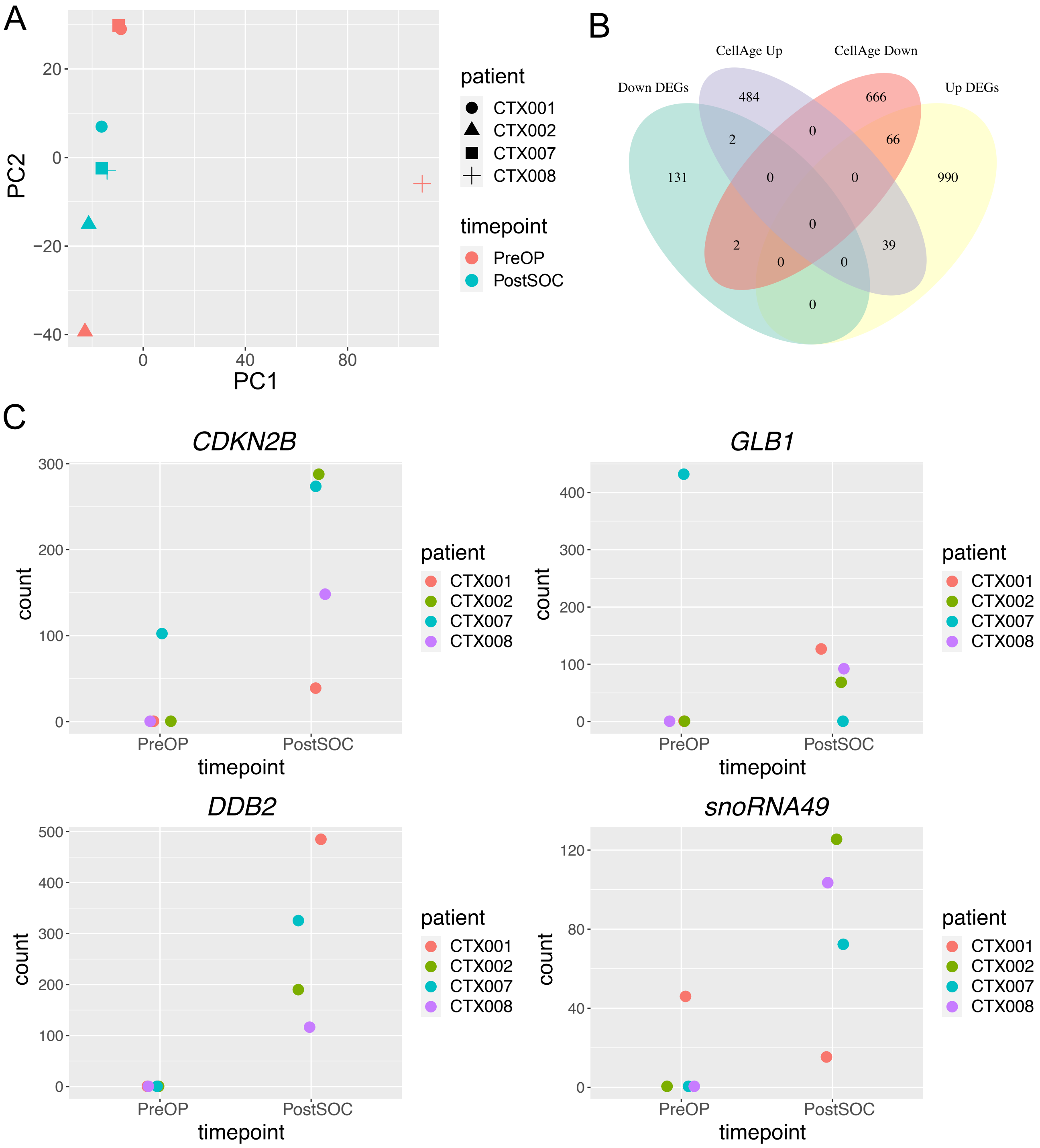

### Figure 6.pdf

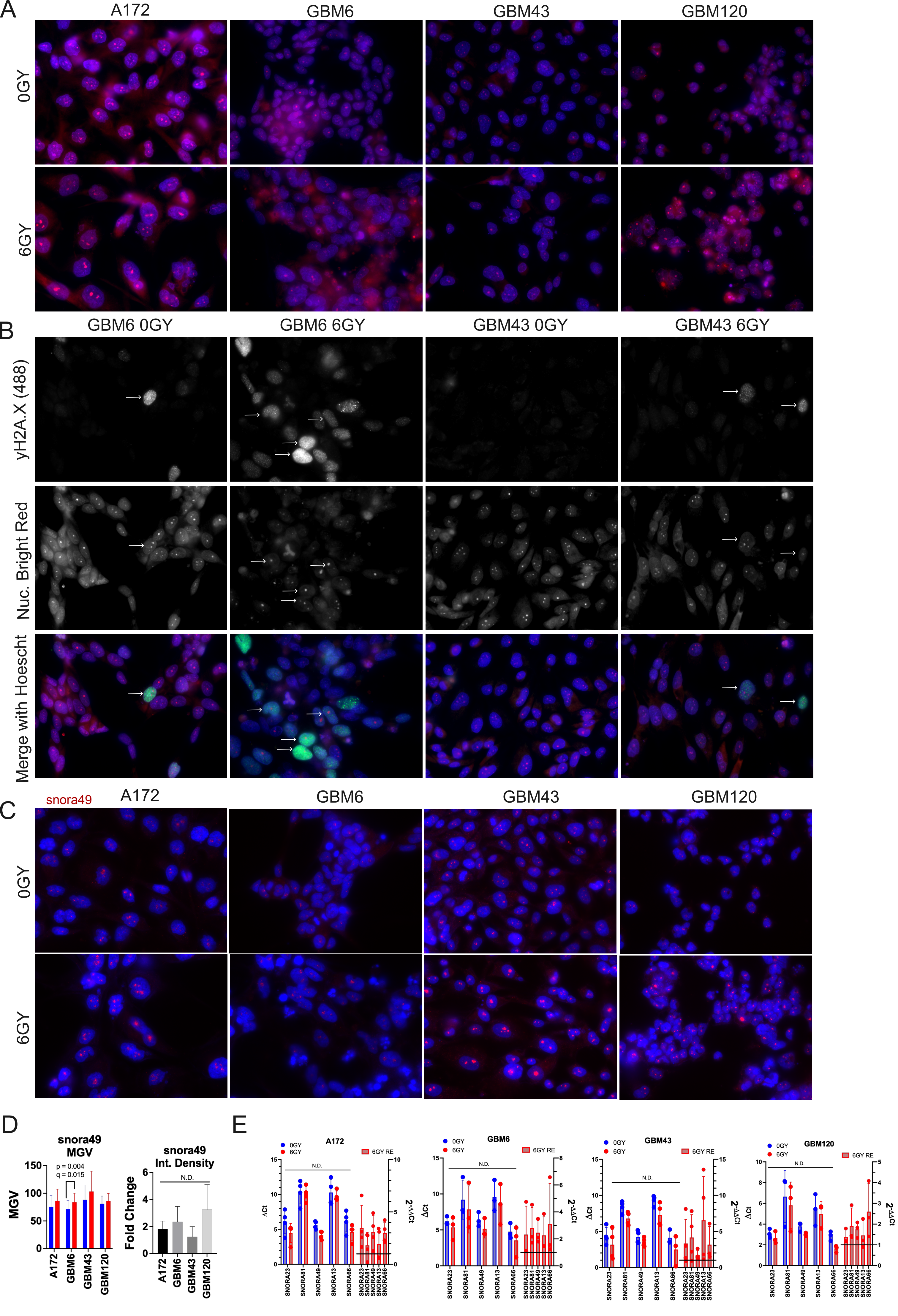

### Supplemental Figure 1.pdf

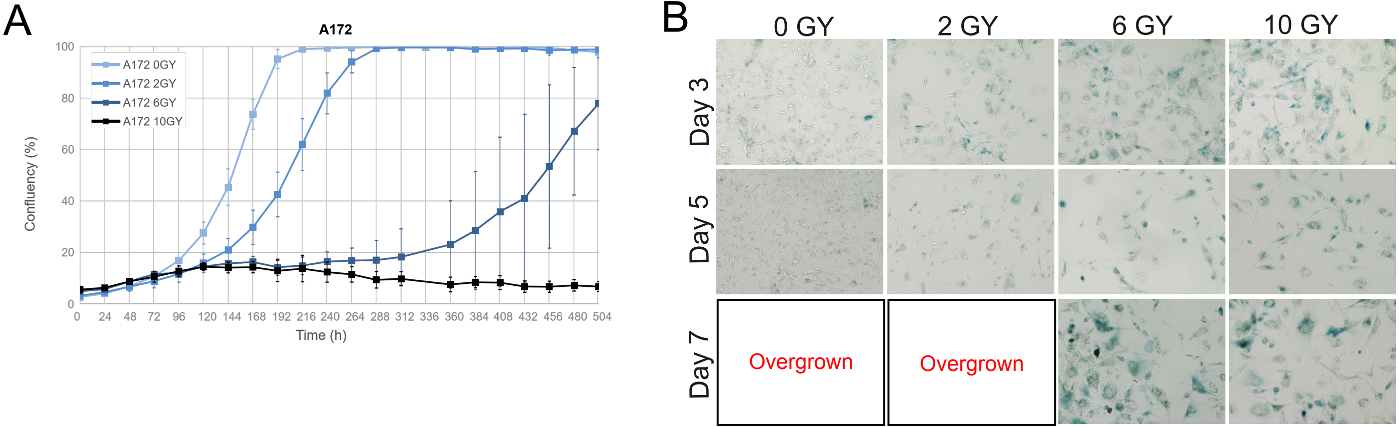

### Supplemental Figure 1.png

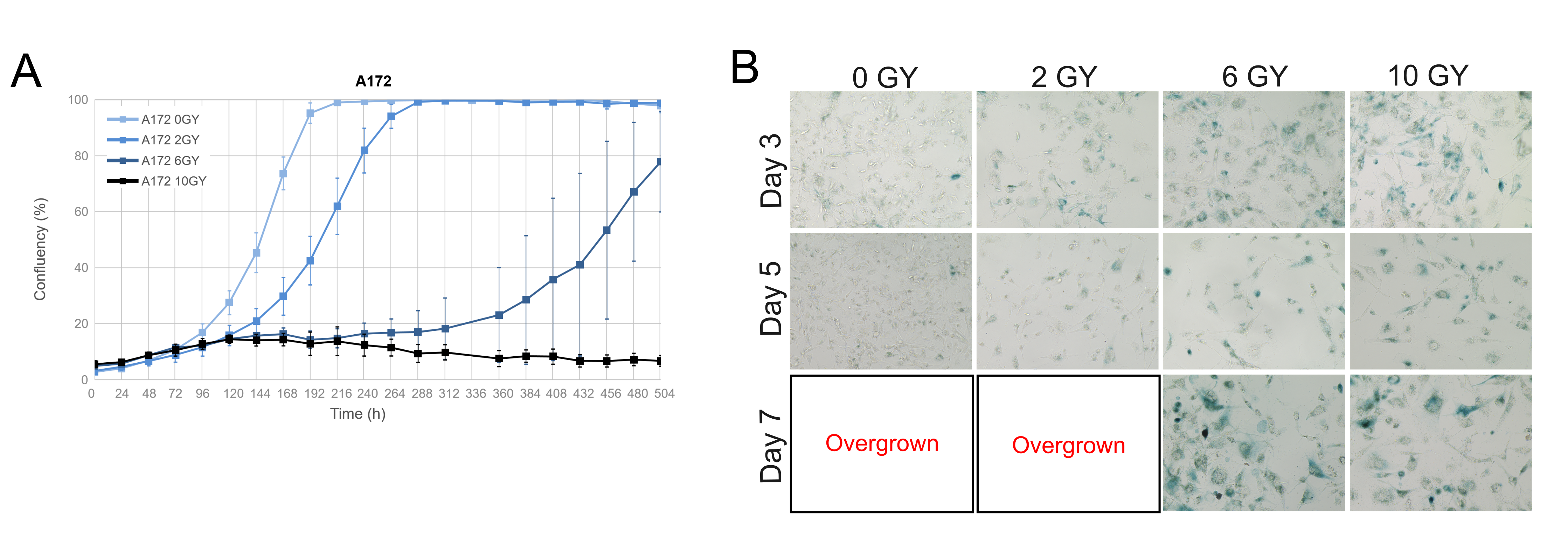

### Supplemental Figure 2.pdf

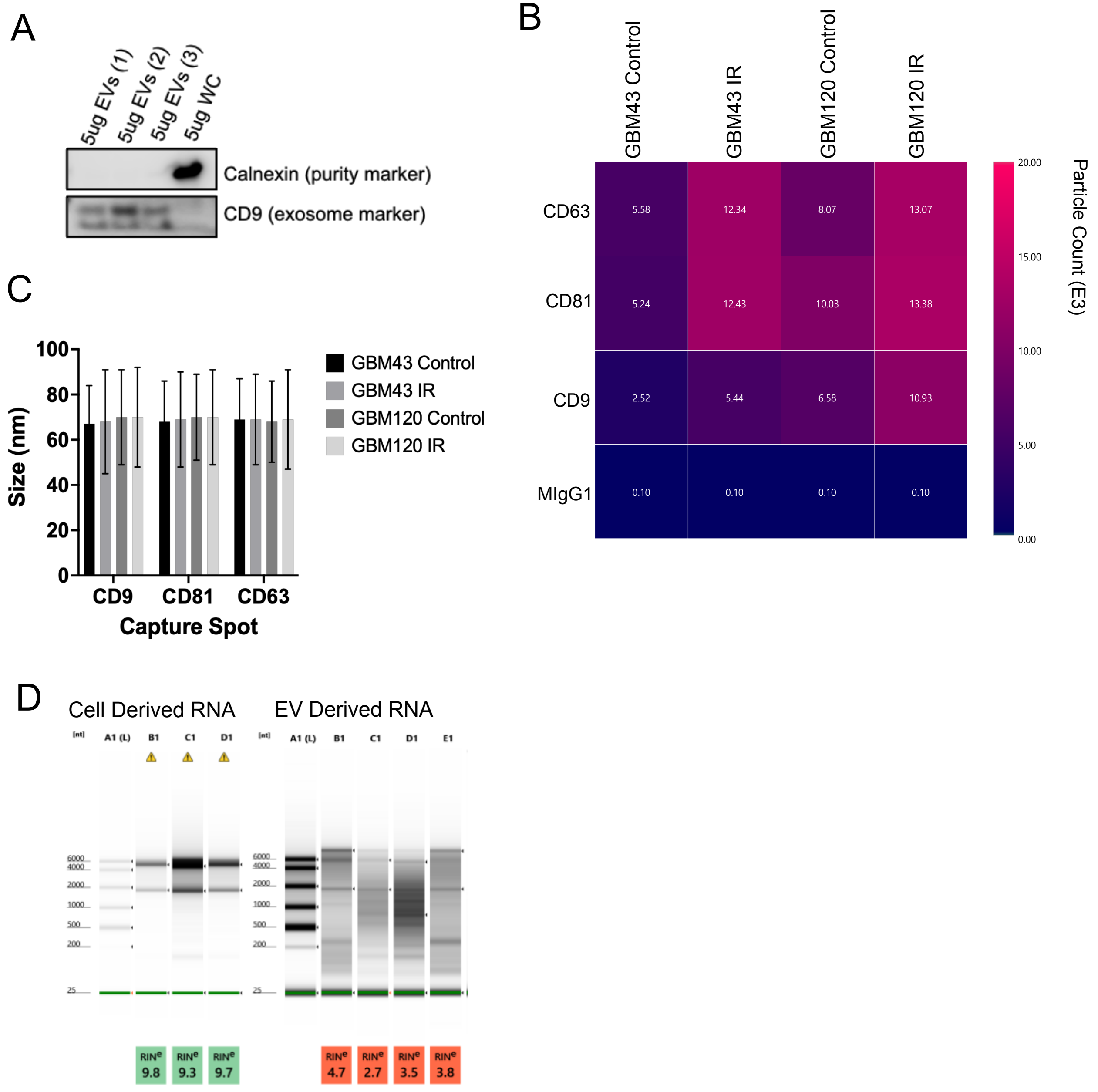

### Supplemental Figure 2.png

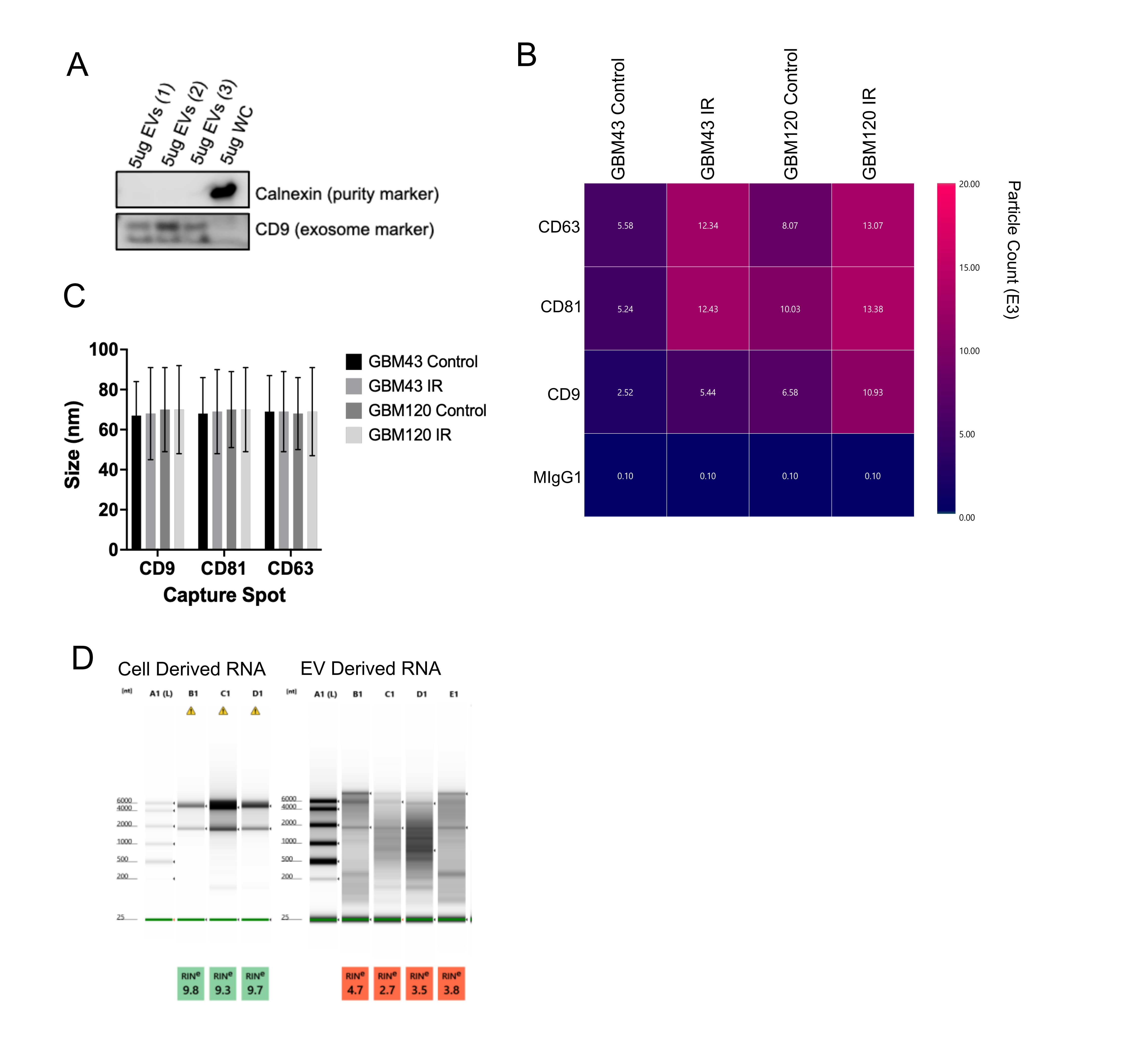

### Supplemental Figure 3.pdf

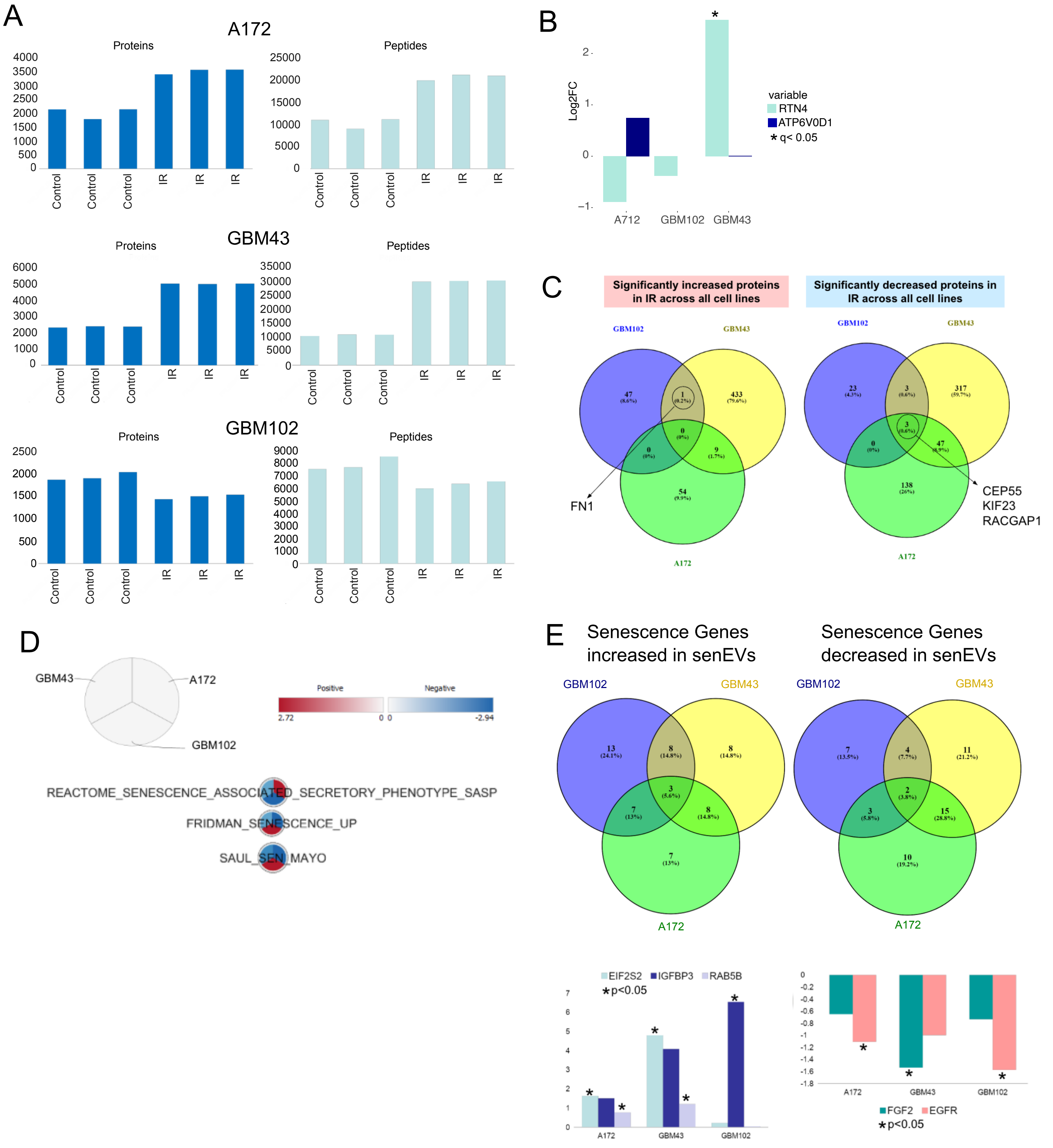

### Supplemental Figure 3.png

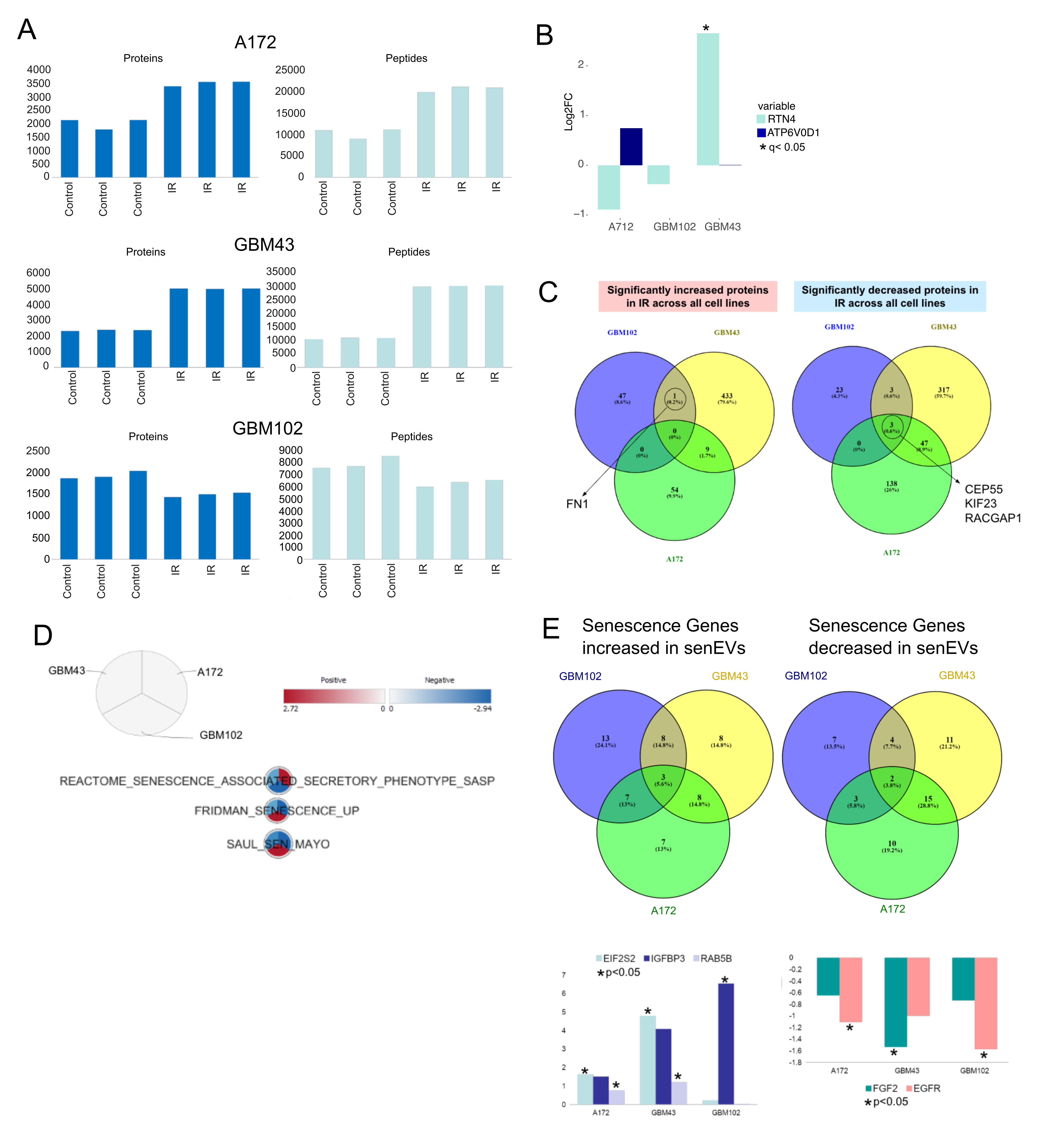

### Supplemental Figure 4.pdf

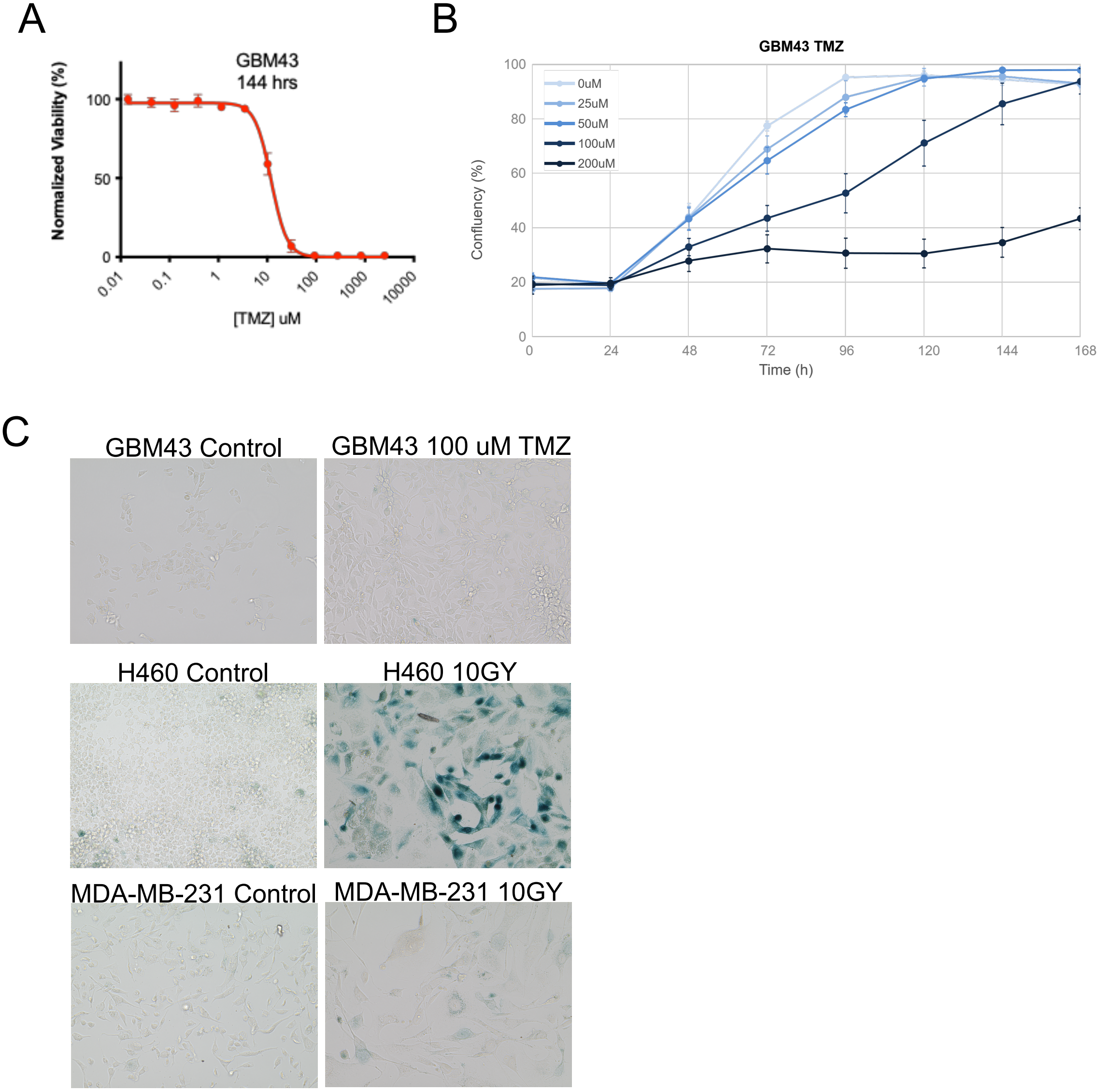

### Supplemental Figure 4.png

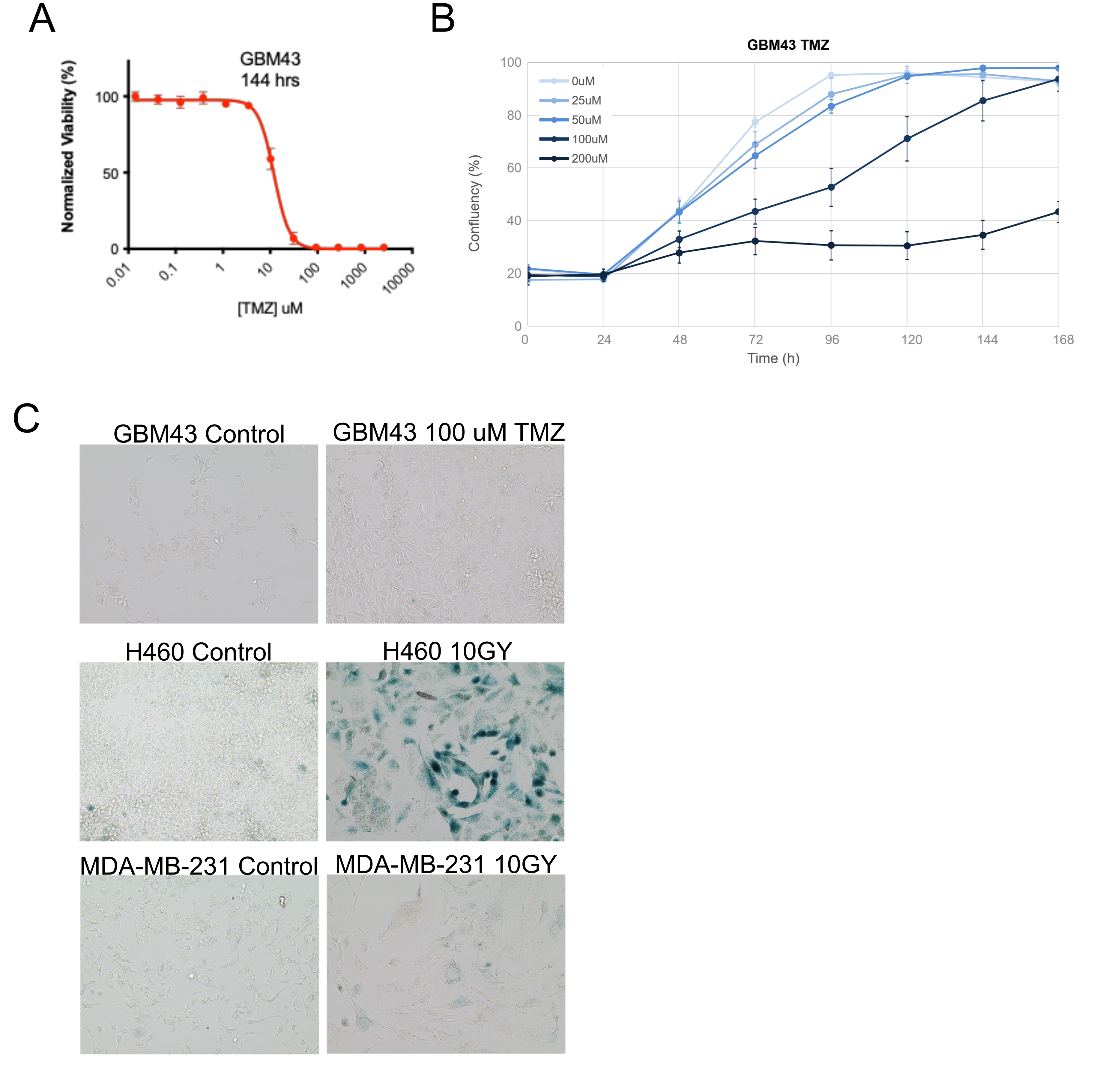

### Supplemental Figure 5.pdf

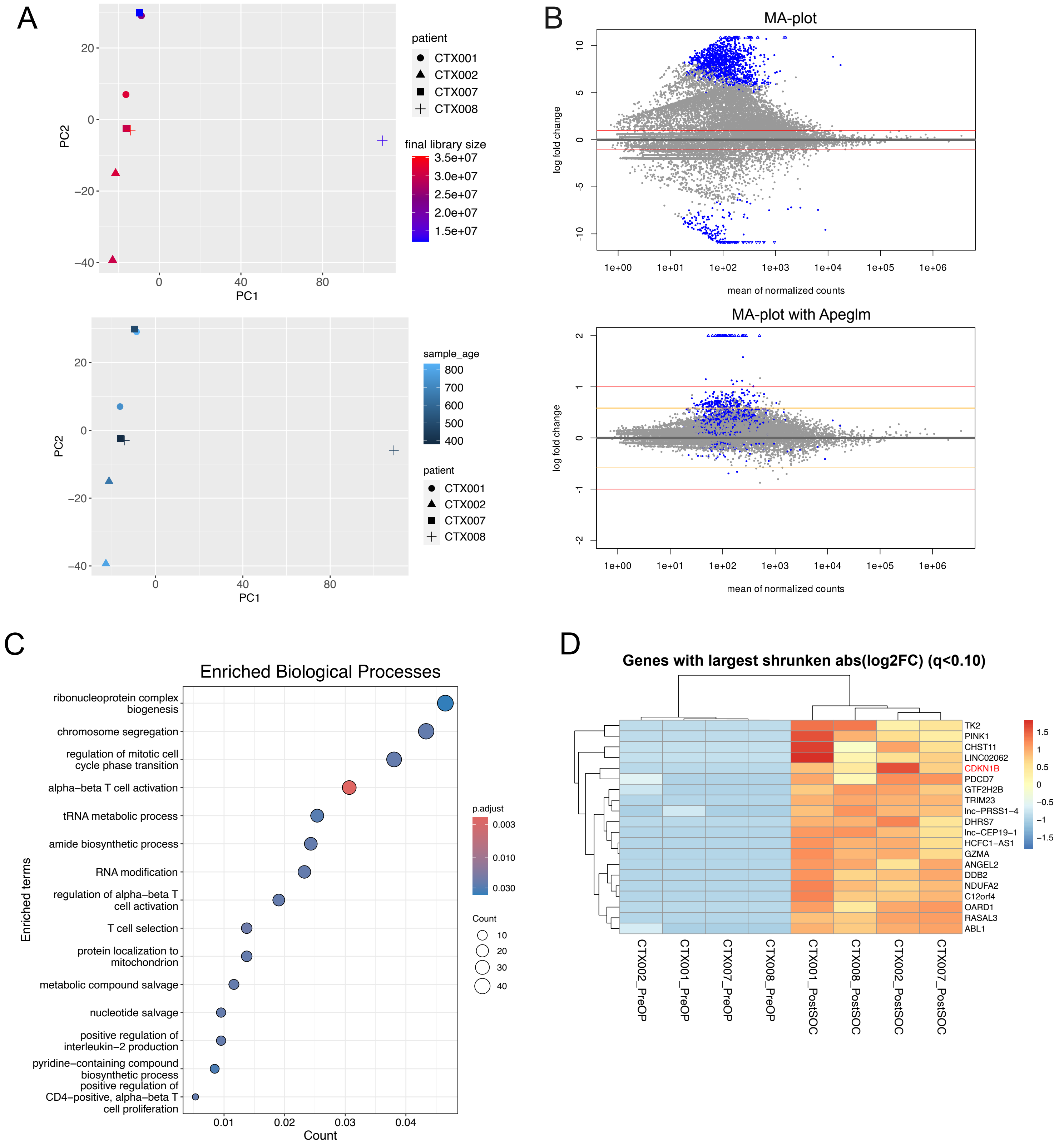

### Supplemental Figure 5.png

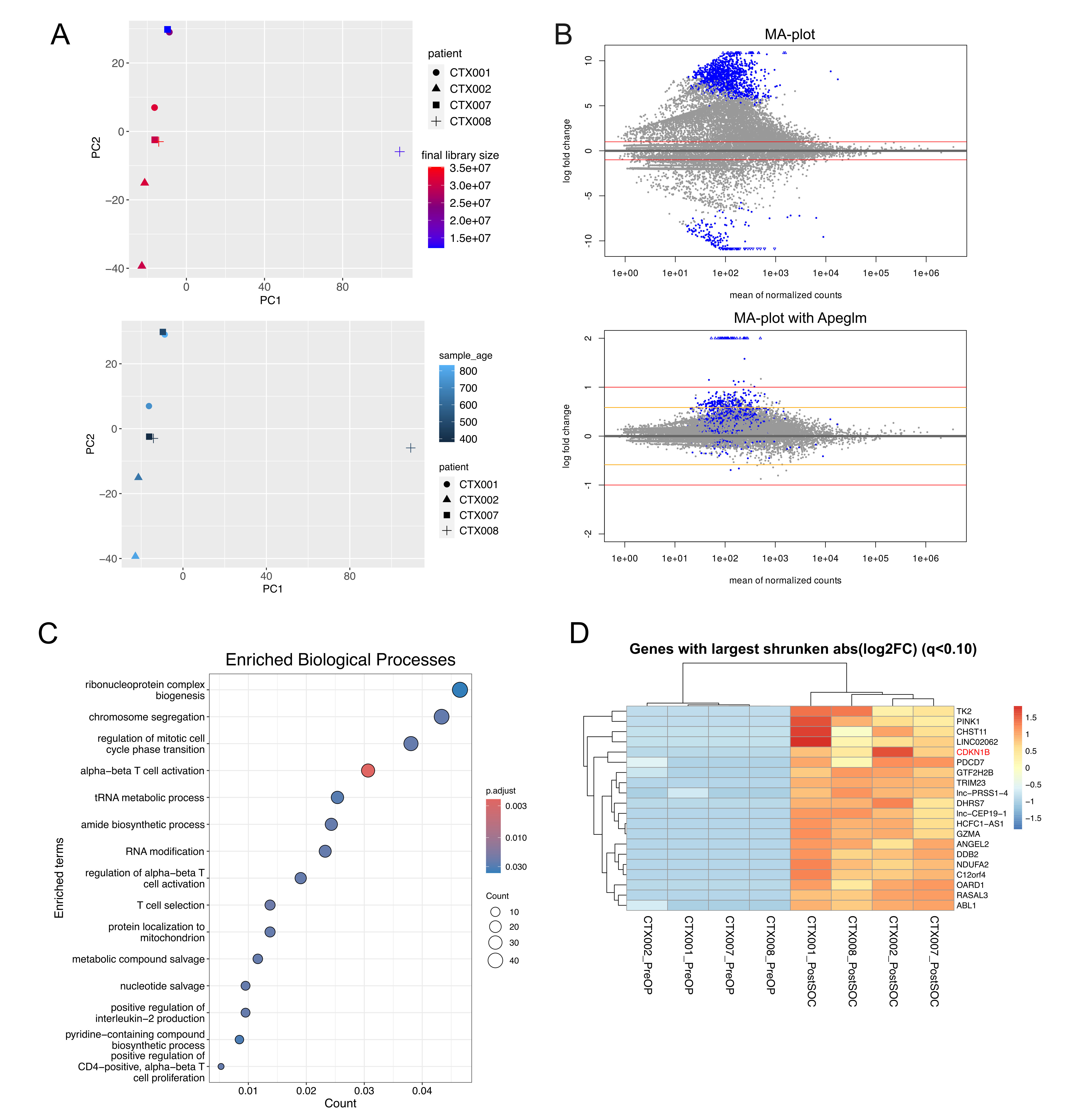

### Supplemental Figure 6.pdf

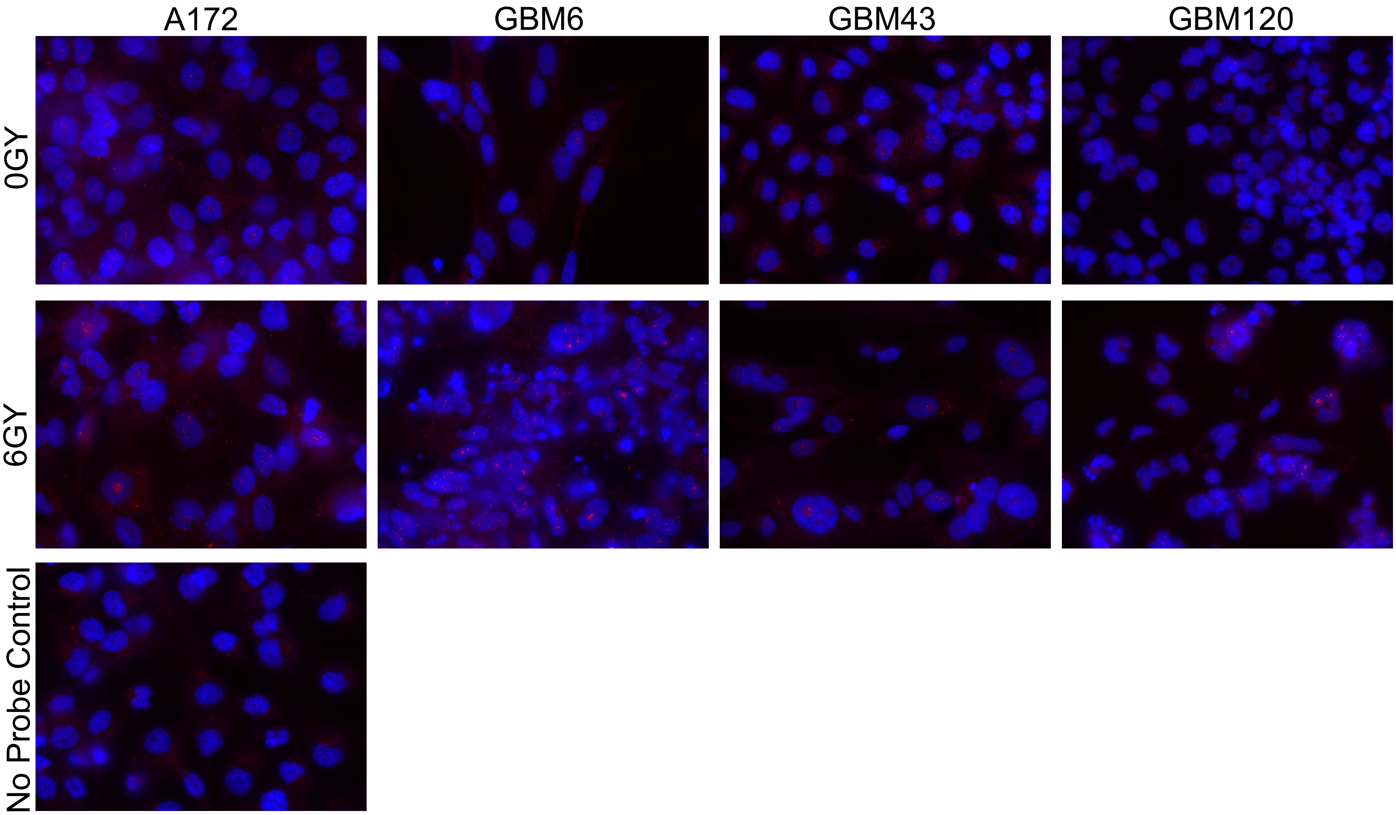

### Supplemental Figure 6.png

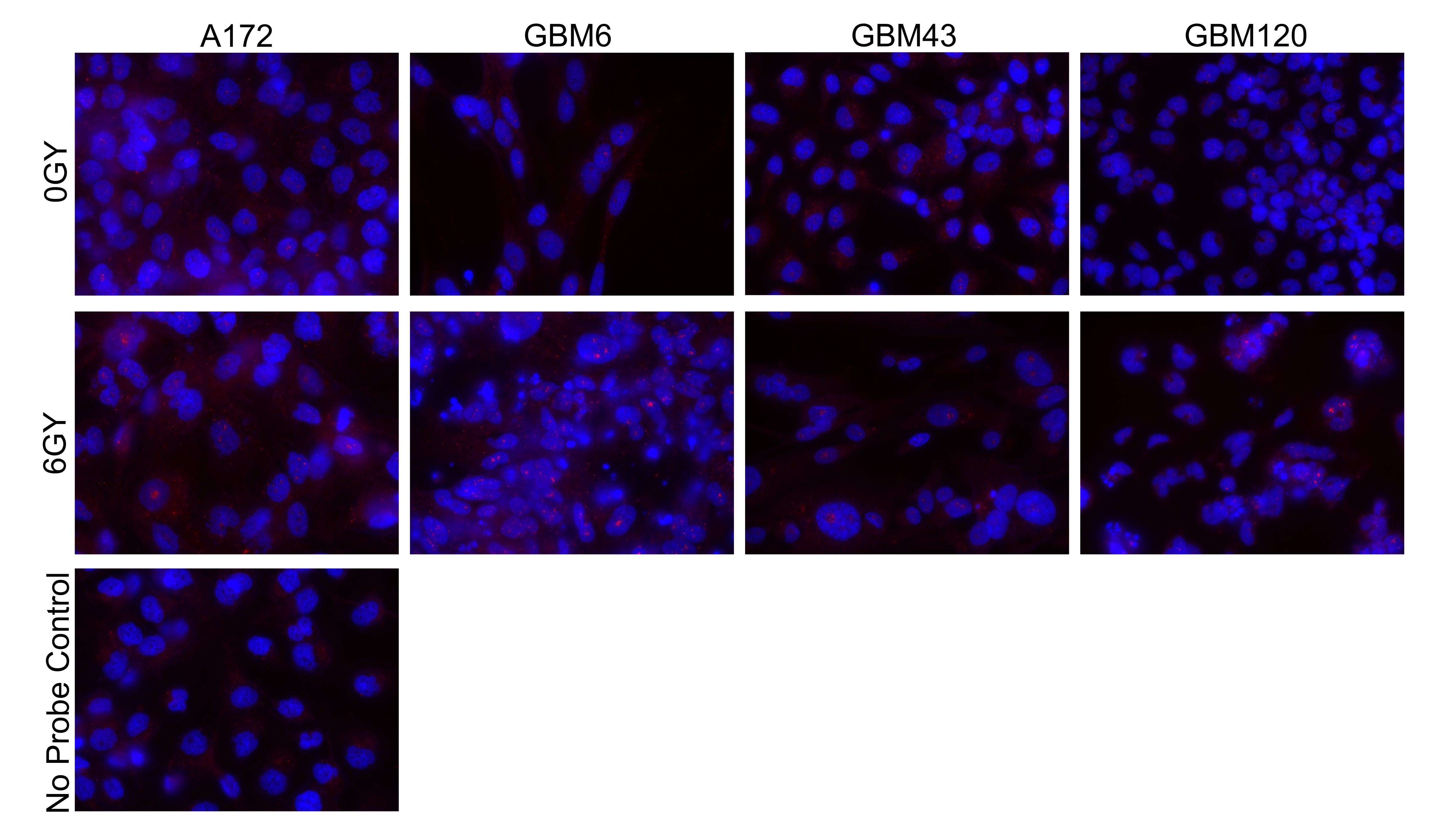

### Supplemental Figure 7.png

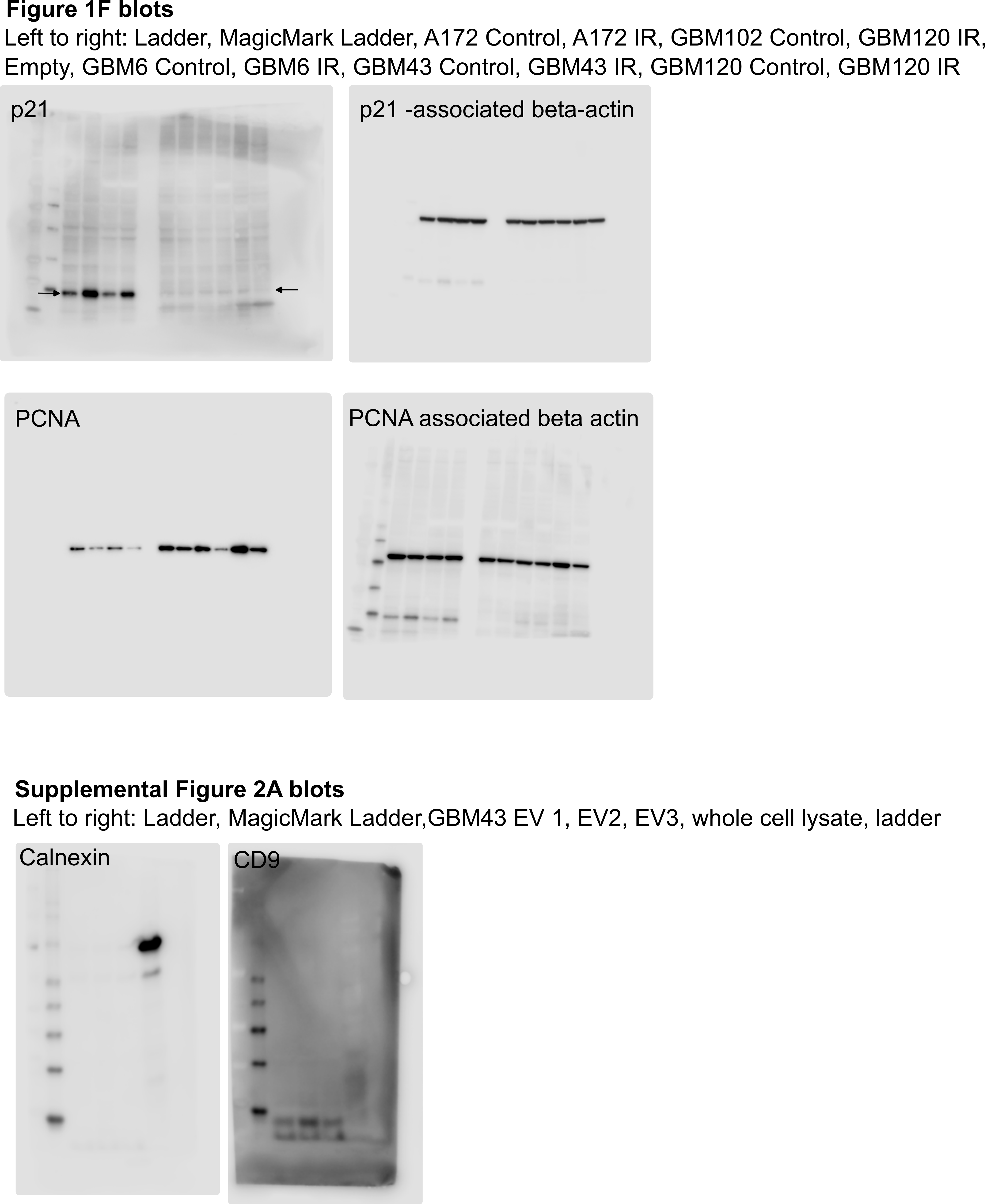
