## Supplementary material for "Molecular profiling of glioblastoma-derived extracellular vesicles identifies small nucleolar RNAs as candidate liquid biomarkers for radiation- induced senescence": High Resolution Images: Supplemental Figure 7.pdf

**Figure 1F blots**

Left to right: Ladder, MagicMark Ladder, A172 Control, A172 IR, GBM102 Control, GBM120 IR, Empty, GBM6 Control, GBM6 IR, GBM43 Control, GBM43 IR, GBM120 Control, GBM120 IR

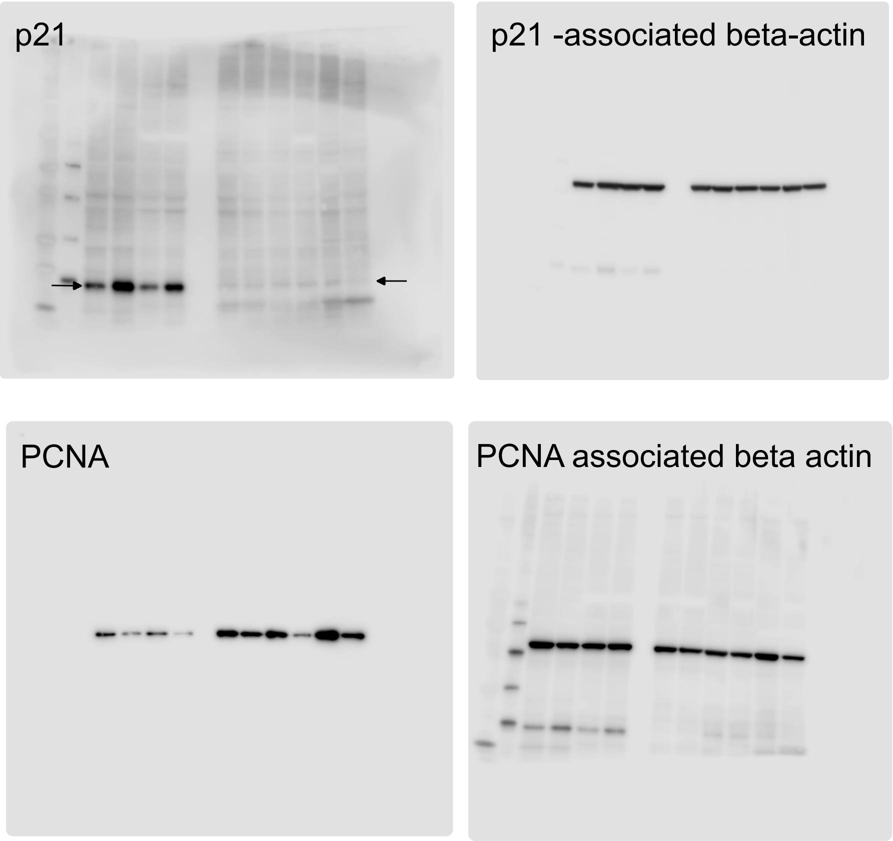


**Supplemental Figure 2A blots**

Left to right: Ladder, MagicMark Ladder,GBM43 EV 1, EV2, EV3, whole cell lysate, ladder

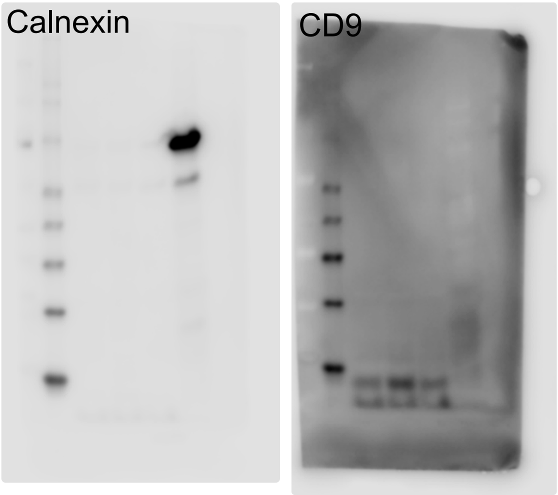
